# Benchmarking active learning protocols for ligand binding affinity prediction

**DOI:** 10.1101/2023.11.24.568570

**Authors:** Rohan Gorantla, Alžbeta Kubincová, Benjamin Suutari, Benjamin P. Cossins, Antonia S. J. S. Mey

**Author notes:** These authors contributed equally to this work (Rohan Gorantla, Alžbeta Kubincová).

## Abstract

Active learning (AL) has become a powerful tool in computational drug discovery, enabling the identification of top binders from vast molecular libraries with reduced costs for relative binding free energy calculations and experiments. To design a robust AL protocol, it is important to understand the influence of AL parameters, as well as the features of the datasets on the outcomes. We use four affinity datasets for different targets (TYK2, USP7, D2R, Mpro) to systematically evaluate the performance of machine learning models (Gaussian Process model, Chemprop), sample selection protocols, as well as the batch size based on metrics describing the overall predictive power of the model (R2, Spearman rank, RMSE) as well as the accurate identification of top 2% / 5% binders (Recall, F1 score). Both models have a comparable Recall of top binders on large datasets, but the GP models surpass Chemprop when training data is sparse. A larger initial batch size, especially on diverse datasets, increased the Recall of both models as well as overall correlation metrics. However, for subsequent cycles, smaller batch sizes of 20 or 30 compounds proved to be desirable. Furthermore, the presence of Gaussian noise to the data, up to a certain threshold, still allowed the model to identify clusters with top-scoring compounds. However, excessive noise (<1*σ*) did impact the model’s predictive and exploitative capabilities.

TOC Graphic

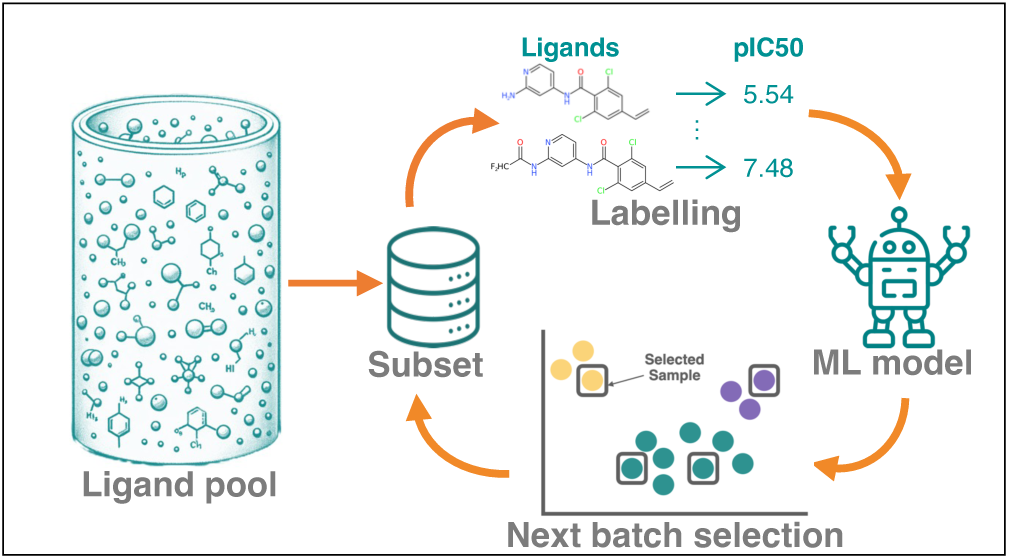

## Introduction

Active learning (AL) is a semi-supervised machine learning (ML) method, which makes use of a model to guide the selection of new samples to label unlabeled data of interest in an iterative process. In the context of computational drug discovery, this method has been used to identify potent inhibitors in small-molecule libraries at a fraction of the cost associated with a systematic potency screen.^1–3^

The identification of drug candidates requires a balance of novelty from exploring new chemical space with optimizing known leads by means of small substitutions. The tension between exploration and exploitation is reflected in AL campaigns, ^2^ and combinations of the two strategies are common, although the exact procedures vary widely.^4,5^ An exploration strategy aims to select samples that are representative of the underlying chemical space in order to construct a good potency model. ^6^ Exploitative strategy, on the other hand, aims to retrieve a high amount of potent compounds by means of a greedy acquisition based on the predicted binding affinity.

Traditionally, AL has been considered in the late stages of lead optimization to select compounds for synthesis. Retrospective studies on affinity data were able to retrieve top binders by using information from a small subset of the data.^4,5^ The procedure was also combined with an automated synthesis setup to select products from a matrix resulting from two types of reagents.^7^ Despite being successful, the throughput in these approaches was low (tens of compounds selected for labeling), and the small sizes of the libraries employed (hundreds of compounds) are often associated with a restricted chemical space.

Computational potency prediction methods have evolved over the last four decades from traditional docking,^8,9^ alchemical free energy (AFE) techniques^10,11^ to more recently machine learning approaches.^12–14^ However, the use of active learning applications together with computational potency estimation such as virtual screening^15–18^ or relative binding free energy (RBFE) calculations using molecular dynamics (MD) simulations^19–23^ only emerged in the past 8 years, driven by the increase in automation and throughput of computational tools for drug discovery. In these cases, 100s-1000 compounds are selected out of pools containing up to 100000 samples. The sheer size of the compound pool goes hand in hand with a high degree of diversity compared to low-throughput use cases, putting more strain on the AL pipeline and necessitates a careful selection of molecular features, ML models and acquisition methods.^15,19,22,23^ In addition to the challenge posed by dataset sizes and diversity, using RBFEs or docking scores in lieu of experimental binding affinities introduces errors of systematic and stochastic nature, which are often not well characterized in advance.

Although active learning presents an opportunity to quickly identify active chemical space in large ligand pools, a routine application of this method in the pharmaceutical industry requires establishing a robust protocol which is transferable between different datasets. Previous active learning studies used RBFE as their labeling tool of choice and only investigated ligands for a single target.^19,21–23^ The scarcity of large public RBFE datasets and the cost and difficulty of generating them is an additional hurdle for establishing robust AL-RBFE benchmarks. Furthermore, none of the studies mentioned above considered cost as a factor in the selection of a protocol, resulting in very large initial batches or exploration phases.^19,22^ They also compared protocols that require a variable amount of RBFE data.^21^ The difference between datasets, their sizes and generation procedures, as well as applied AL protocols, and different combinations of metrics to evaluate the performance of AL makes it difficult to compare literature protocols and identify best-practice approaches.

Here, we evaluate AL protocols in a rigorous manner by using four publicly available datasets for benchmarking. These datasets differ in their protein targets, kind of potency measurement (Δ*G* from RBFE or experimental *K_i_* / IC50), size (600 to 10000 samples), and degree of diversity. Furthermore, we use a wide range of metrics to gain a holistic perspective, ranging from conventional regression metrics such as R2 to assess the overall performance of the ML model, to Recall and F1 scores for the top 2%/5% binders to assess the exploitative capabilities of a model and the degree of exhaustion of active chemical space. To investigate the benefit of pre-trained model architectures for AL, we compare a finetuned Chemprop model with Gaussian process (GP) regression, which is a common choice for AL. ^21,22^ Finally, the total number of acquired samples is always kept constant throughout all experiments to compare AL protocols at a fixed cost. We leave the choice of how unlabeled data is labeled up to the user and could be a docking score, experimental value, or RBFE calculation. However, mixing these may require care in accounting for accuracy or trustworthiness of the labeling method used.

The paper is structured as follows. In the Methods section, we give a detailed overview of the datasets and provide details of the AL procedure, ML models, and metrics. In the Results section, the GP and Chemprop models are first benchmarked to assess differences in their predictive power between datasets. Next, we compare AL procedures by varying the size of the initial batch and the method for its acquisition. Thereafter, we identify an optimal batch size for cycles that come after the initial batch(es). Finally, we assess the robustness of AL towards stochastic noise in the potency data.

## Methods

### Datasets

We used four publicly available binding affinity datasets that encompass the following protein targets: Tyrosine Kinase 2 (TYK2); a G protein-coupled receptors target, Dopamine Receptor D2 (D2R); and two proteases, Ubiquitin-Specific Protease 7 (USP7) and SARS-CoV-2 Main Protease (Mpro). Table 1 provides the number of ligands in each datasets as well as other information relevant for subsequent AL experiments. Fig. 1A shows the distribution of the measured affinities associated with each dataset. All the datasets used in our study are accessible at https://github.com/meyresearch/ActiveLearning_BindingAffinity.

**Table 1:** Summary of dataset characteristics for protein targets used in our study. The table presents the total number of ligands, the binding measure used for training and inference, the percentage of data utilized for AL based on a consistent sample of 360 compounds acquired over AL cycles, and the count of compounds in the top 5% and top 2% fraction of the dataset.

| Target | Ligands | Binding measure | % Data for AL | Top 5% | Top 2% |
| --- | --- | --- | --- | --- | --- |
| TYK2 | 9997 | $\text{pK}_i$ | 3.6 | 500 | 200 |
| USP7 | 4535 | $\text{pIC}_{50}$ | 7.9 | 227 | 90 |
| D2R | 2502 | $\text{pK}_i$ | 14.4 | 125 | 50 |
| Mpro | 665 | $\text{pIC}_{50}$ | 54.1 | 33 | 13 |

The **TYK2** dataset was derived from the work of Thompson et al., ^21^ which focused on optimizing the AL methodologies for RBFE calculations. This dataset comprises of 10000 congeneric molecules targeting the TYK2 kinase, all of which were synthesized using an amino-pyrimidine core scaffold. The dataset was initially populated with 573 TYK2 inhibitors, which were subsequently decomposed into unique R-groups at three attachment points. These groups were then combinatorially assembled to create an initial library of 203406 unique compounds, which was then filtered down to 10000 molecules based on a set of “drug-like” properties. ΔΔ*G* values obtained from RBFE calculations were converted to pK*_i_* values, providing a more interpretable metric for binding affinity, using

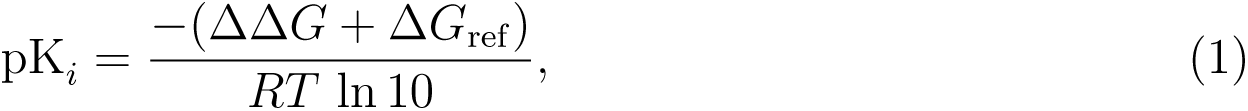

where pK*_i_* is the negative logarithm of the inhibition constant, which is a measure of the binding affinity of a ligand for its target (TYK2). ΔΔ*G* is the free energy difference of binding between a reference compound and a second compound. Δ*G*_ref_ is the absolute binding free energy of the reference compound (-47.778 kJ/mol for TYK2), *R* is the universal gas constant, approximately equal to 8.314 J/(mol·K), and *T* is the absolute temperature. In Fig. 1B, the Uniform Manifold Approximation and Projection (UMAP)^24^ of the TYK2 dataset shows clear clusters that capture variations in R-groups attached to the core scaffold. We can see that most of the active compounds are located within the two upper clusters. Fig. S1 in the SI highlights that the majority of the top 2% binders are situated here.

The **USP7** dataset was curated by Shen et al., ^25^ with the primary objective of building a classification model to distinguish active from inactive inhibitors. The SMILES of over 4000 ligands together with their experimental affinities including *K*_i_, *K*_d_ and IC50, were collected from ChEMBL.^26^ Duplicate SMILES were aggregated into unique entries using the Open Babel package 2.3.1^27^ by Shen et al.^25^ All experimental results with varying units were converted to IC50 values for each SMILES, which, for the scope of our study, were translated to pIC50 values. As evident from Fig. 1A, the USP7 dataset exhibits multiple assay minima. Moreover, the UMAP visualization with cluster centroids in Fig. 1D highlights the dataset’s diverse core scaffolds and R-groups, showing the presence of heterogeneity. However, scaffolds tend to be well-preserved within a cluster. A significant portion of the top active compounds cluster in the upper regions of the UMAPs.

The **D2R** dataset is a subset of the ACNet dataset, ^28^ which was curated from the ChEMBL^26^ database (version 28) on 190 targets to study the performance of machine learning models on data from activity cliffs. Zhang et al.^28^ categorised matched molecular pairs as activity cliff if Δ in potency is pK*_i_ ≥* 2. We selected the D2R target due to a high number of associated activity data, making it particularly suitable for our study. The ACNet dataset was constructed by screening over 17 million activities, only retaining compounds tested against single human targets in direct interaction binding assays and filtering data with low assay confidence. The dataset uses assay-independent equilibrium constants (pK*_i_*) as the measure of potency. We retained the pK*_i_* values and averaged over duplicate entries with the same SMILES to ensure data consistency, leading to a reduction from 4121 to 2502 entries. From the UMAP in Fig. 1D, the D2R dataset appears to encompass a heterogeneous assortment of compounds, with a large number of congeneric series of 10-20 compounds each. The absence of distinct clusters and the dispersed distribution of binding scores suggest intricate structure-activity relationships.

The **Mpro** dataset is part of the COVID Moonshot project,^29^ which focuses on the development of inhibitors for the SARS-CoV-2 main protease. The dataset provides experimental pIC50 values, which are an amalgamation derived from single enantiomers and racemic mixtures. The project consists of several design cycles (sprints) with medicinal chemists selecting compounds for synthesis out of a database with public submissions. The UMAP depicted in Fig. 1E, shows a diverse composition of the Mpro dataset. The absence of distinct clusters and dispersed distribution of top active compounds highlight its intricate nature. Importantly, with only 665 compounds, this dataset is significantly smaller than the others in our study, offering a distinct context for AL pipeline investigations in low-data settings.

**Figure 1:**
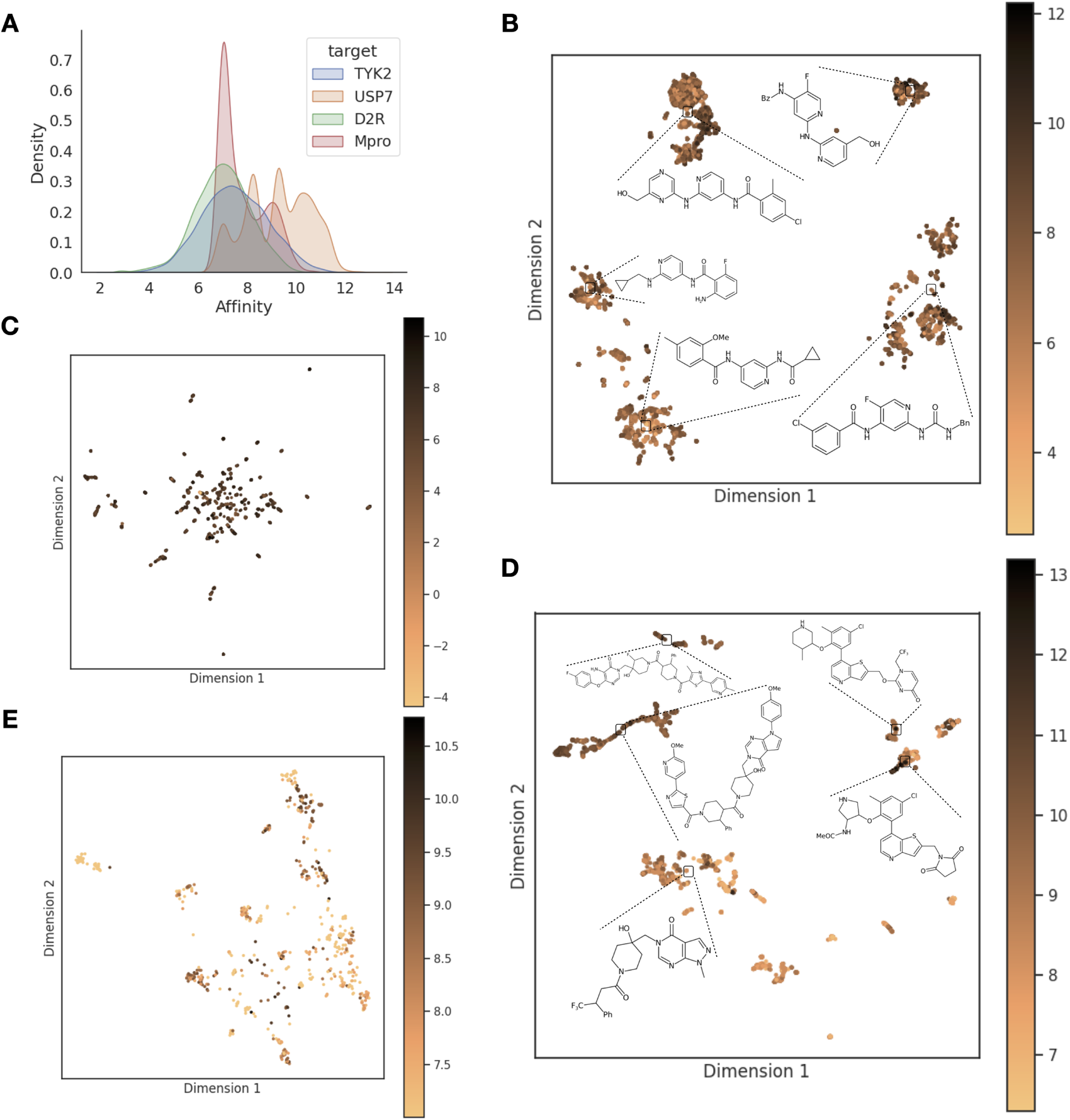
Distribution of affinity scores and UMAP projections for four protein targets. **A:** Kernel Density Estimation plot illustrating the distribution of affinity scores for each target dataset: pK*_i_* values for TYK2 and D2R, and pIC50 values for USP7 and Mpro. **B:** UMAP projection of the TYK2 dataset with overlaid cluster centroid compounds. **C:** UMAP visualization of the D2R dataset. **D:** UMAP projection of the USP7 dataset with overlaid cluster centroid compounds. **E** UMAP representation of the Mpro dataset.

### Active learning (AL) protocols

AL is a machine learning paradigm designed to optimize the selection of samples for training models (see Fig. 2). It is particularly useful for problems where labels can be calculated for every data point, but the associated computational cost is high, as is the case for RBFE calculations or testing the data point in the lab. The key difference from conventional machine learning consists of splitting the training process into several AL cycles such that a model trained on a subset of samples informs the selection of the next batch to be added to its training set.

**Figure 2:**
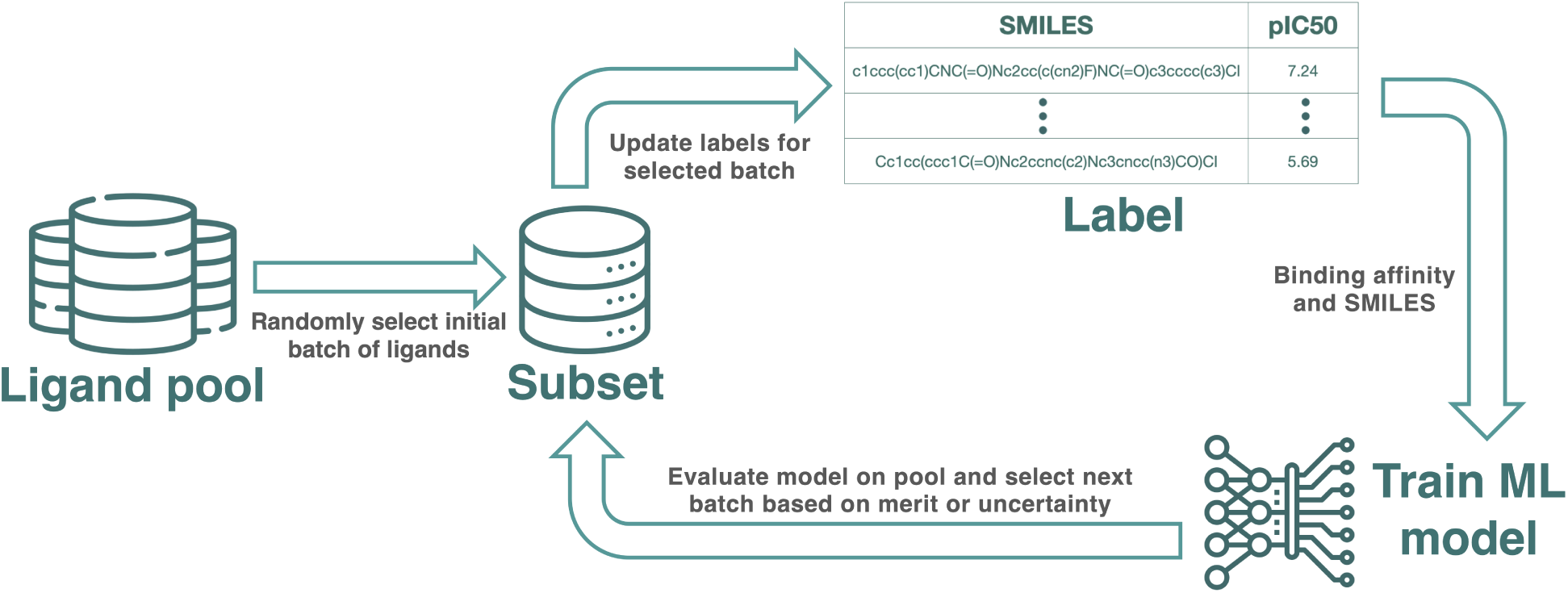
Schematic overview of the AL pipeline. An AL cycle begins with a randomly chosen batch from the available pool of compounds, followed by labeling and model training on the subset with labels. The subsequent batch for the next AL cycle is strategically chosen based on model predictions and uncertainties using exploration, exploitation, or random strategies.

The initial batch of compounds was selected at random, as this is a common strategy in previously published studies. ^15,21^ This choice also ensures that the distribution of training data matches the pool used for inference, which is not the case in diversity-based selection methods.

In a real AL use case, labels for the selected compounds can be acquired by means of RBFE calculations. We take advantage of experimental potency values from the literature instead of RBFE calculations. This provides a robust insight into AL on retrospective data.

A model is then trained on a subset of labeled samples and used to make predictions on unlabeled data. On the basis of these predictions, a strategic subset of samples is selected for labeling, often employing strategies aimed at either “exploration” of the chemical space or “exploitation” of promising regions within it. These newly labeled samples are then incorporated into the training set for the next AL cycle.

In each AL cycle, we employ one of the strategies for sample selection: random, exploration, and exploitation. Random selection involves choosing compounds arbitrarily from the remaining data. Exploitation focuses on selecting compounds with the highest predicted potency, thereby exhausting high-potency areas in the chemical space. Exploration selects compounds with the highest prediction uncertainty, aiming to sample broadly across the chemical space to gain a more global understanding, thereby potentially identifying new promising areas in chemical space.

To ensure a fair evaluation for different targets, we always acquire a total of 360 compounds for labeling over the whole AL procedure, irrespective of the dataset, model, or selection protocol. The number of AL cycles is always adjusted to fit this total, depending on the batch sizes. Additionally, to account for variability, each experiment was conducted three times using different seeds for the initial batch selection.

### ML models

#### Gaussian Process Regression

We chose Gaussian Process (GP) regression^30^ for its ability to provide both expected values and uncertainty estimates for predictions, as well as its proven performance in a previous AL study on the TYK2 target by Thompson et al. ^21^ GP is a non-parametric Bayesian approach that provides a probabilistic framework for making predictions. GPs make use of a kernel function to measure the similarity between data points. The kernel function is used to construct a covariance matrix of observed features, which is used to make predictions for unseen data. In our implementation, we use the Tanimoto similarity kernel, which is particularly useful for measuring the similarity between sets, making it apt for binary fingerprints. Our implementation of GP is based on the GPyTorch version 1.10^31^ library.

By default, ligands were featurized using ECFP8 fingerprints from OpenEye’s OEChem toolkit version 3.4.0.0. ^32^ To assess the influence of chiral descriptors on GP model performance, an alternative featurization using Morgan fingerprints with chirality descriptors was also assessed as a part of benchmarking. Morgan fingerprints were generated using RDkit, ^33^ and have a radius of 4. Both ECFP8 and Morgan fingerprints were hashed to 4096 bits.

#### Chemprop

Chemprop (CP) is a message-passing neural network designed for molecular property prediction. The model operates by iteratively updating the atom and bond features of a molecule through message-passing layers (and hence does not rely on molecular fingerprints such as GP). This allows the model to capture both local and global structural information. Chemprop has been shown to perform well on a variety of molecular property prediction tasks, including solubility, toxicity, and binding affinity.^34^ Monte Carlo dropout^35^ was used to provide a measure of uncertainty. Our implementation is based on the Chemprop package version 1.6.1, ^36^ and we use a model pre-trained on potency data across 1788 targets, including Kinases, GPCRs, and Proteases. The data for pretraining the model is mostly taken from ChEMBL^26^ for targets having more than 200 interaction data points. For our work, we unfreeze the entire encoder and train the model in each AL cycle for 500 epochs with a batch size of 50 and a learning rate of 0.0001 for 10 warmup epochs, ramping up to a maximum of 0.001 for the remaining epochs using Noam scheduler. These hyperparameters for fine-tuning the Chemprop model are empirically determined.

### Analysis

We used a range of metrics to evaluate the performance of our AL benchmark. These metrics are selected to assess both the exploitative and predictive aspects of the model. Whereas exploitation is governed by the predictive prowess of the model on the high end of the potency range, regression metrics reflect its performance on the bulk of the data.

To define Recall and F1 score in this context, the selection process is converted to a classification task. A compound is considered “True” if it has been acquired for labeling in any of the previous AL cycles, and “False” otherwise. We further categorize compounds into “active” and “inactive” based on their relative ordering, specifically focusing on the top 2% and top 5% of compounds. The Recall metric is calculated according to Equation 2,

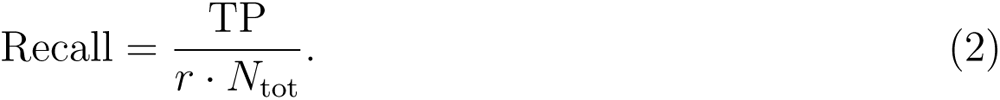

Here, TP represents the number of true positives, *r* is the fraction of compounds considered true (either 0.02 for the top 2% or 0.05 for the top 5%), and *N*_tot_ is the total number of samples in the given dataset. The expectation value of TP for a random selection is given by *r · N*_acq_, where *N*_acq_ represents the number of compounds acquired or selected in the current AL cycle. The F1-score is particularly useful for assessing the models ability to correctly identify the most promising compounds (TP) while minimizing the selection of false positives (FP). We compute the F1-score according to Equation 3,

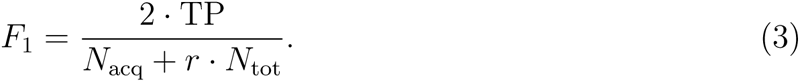

Here, we used the relations *N*_acq_ = TP + FP and *r · N*_tot_ = TP + FN. False negatives (FN) represent the promising compounds that the model failed to identify.

To assess the predictive power of the models, we use the coefficient of determination R2, Spearman *ρ*, and the Root Mean Square Error (RMSE). To visualize the high-dimensional fingerprints on two-dimensional maps, we use Uniform Manifold Approximation and Projection (UMAP)^24^ as a dimensionality reduction technique using the umap-learn package version 0.5.3.^37^ In contrast to widely used tSNE plots, UMAPs do not rely on a fixed cutoff and instead keep the number of neighbors constant (here 50), which is an arguably better choice for preserving the global structure of the data.

## Results and discussion

### Model benchmarking

Fig. 3 compares GP and CP models trained on a 5-fold split, where the training set was made up of 20% of the data, and metrics were calculated on the test sets containing the remaining 80%. Error bars represent a 95% confidence interval across these folds. The calculated metrics are the coefficient of determination R2 (A), Spearman *ρ* (B), as well as, the Recall of top 2% (C) and 5% (D) samples across the test sets. To evaluate the impact of chiral molecules, we considered a GP model trained on Morgan fingerprints with chirality descriptors in addition to achiral ECFP8 fingerprints with the same size and radius.

**Figure 3:**
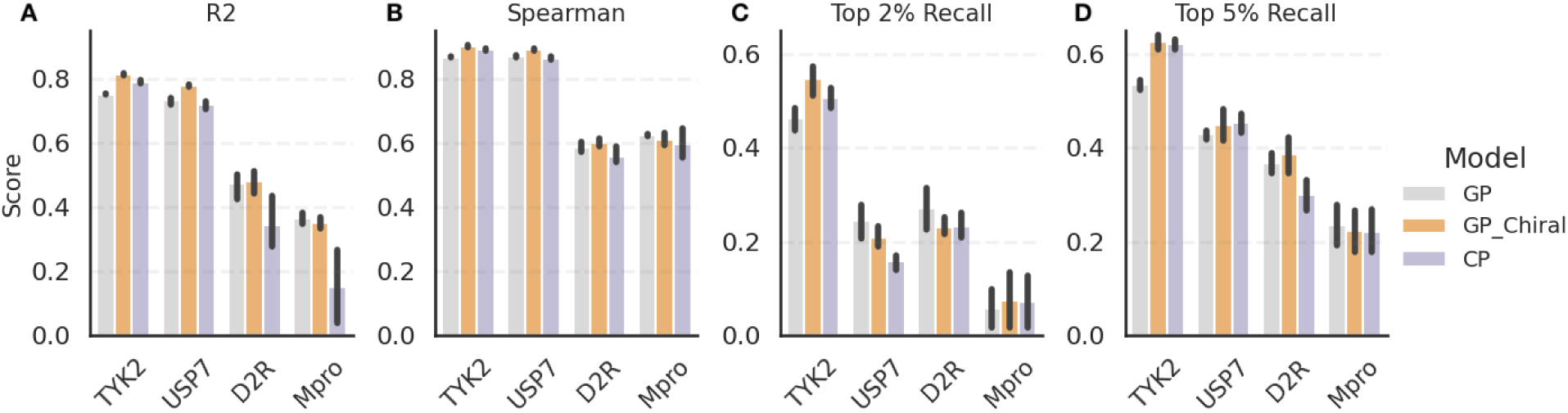
Benchmarking GP and CP model on four target datasets (TYK2, USP7, D2R, Mpro) used in our AL study. We use 20% of the dataset for training the models, and the remaining 80% as a test set to calculate the metrics-**A:** R2, **B:** Spearman *ρ*, and **C:** Recall for top 2% compounds and **D:** Recall for top 5% compounds. Datasets are sorted based on their size i.e., from large to small (see also Table 1). While both models exhibit robust predictive power, CP is more sensitive to the dataset size than GP, and the introduction of chirality descriptors offers limited enhancement in model accuracy.

The goal of this work is to assess the performance of the two models on each dataset in the limit of a large training set. However, the training sets amount to a fraction of the pool rather than being comparable in size to reflect the total number of compounds typically acquired in AL. Table 1 shows that the subsequent AL experiments acquire less than 20% of the data in total except for Mpro, where over half of the data is acquired. Given the lack of bias in a random sample selection, this analysis provides an upper bound for the model performance in terms of regression. It also gives an estimate for the Recall given a large amount of training data.

For each dataset, all of the models demonstrated predictive capabilities, with an R2 larger than 0.3 (with the exception of CP trained on Mpro data), Spearman *ρ* over 0.5 and top 5% Recalls of 0.2 or more. This suggests that it is possible to train a predictive model even on the more challenging datasets given enough training data sampled broadly over the available chemical space. There is, however, a clear trend in model performance concerning the dataset size. R2 and top 5% Recall monotonously decrease with decreasing size of the datsaset. The trend is also present, to a weaker extent, for Spearman *ρ* and the top 2% Recall, which is more pronounced for CP compared to GP. This observation aligns with the understanding that CP, being a deep learning model, benefits from larger datasets.

Both Mpro and D2R datasets presented challenges due to their heterogeneous nature. The lack of distinct clustering in the UMAP projections (Fig. 1C, E) for these datasets, especially for D2R, indicates the diverse composition of compounds, making them potentially harder to fit with machine learning models. Spearman *ρ* for these two datasets is indeed lower than for TYK2 and USP7. However, the comparably large training set accounts for diversity to some degree, which is why differences between different types of datasets are not very pronounced in this benchmark.

The GP model using chirality descriptors showed comparable performance to the model trained on achiral fingerprints, suggesting that introducing chirality representation did not significantly enhance the models performance. For Mpro and D2R compounds, which have a significant amount of chiral centers, the performance remained comparable between both the fingerprints. Similarly, the USP7 dataset, which contains only a few chiral compounds, showed no significant improvement with the chirality descriptors. It is likely that chirality does not play an important role in the structure-activity relationship of these series, or that stereochemistry information is missing for some of the data (such as TYK2).

### Active learning strategies

In this section, we investigate the impact of batch sizes, sample acquisition strategies, as well as the exploration-exploitation trade-off in simulated AL scenarios. The focus herein lies on protocols with a distinct exploration phase, which aims at selecting diverse samples for building a robust and predictive model, followed by greedy acquisition using the merit predicted by the model (exploitation). Splitting the AL protocol in these two stages simplifies the evaluation of the individual benefits of each phase. Note, however, that the interleaving of the two strategies has also been studied in the literature^5,20,23^ yielding good results. We always acquire a total of 360 compounds for labeling to fairly evaluate selection protocols that differ in their batch sizes.

#### Selection of initial samples

In an AL setting, it is essential to understand the influence of the initial sample selection strategy as it guides the trajectory of subsequent AL cycles. We explore initial batch sizes and strategies for the selection of these compounds using three distinct selection protocols. All of them are initiated with 60 samples selected at random (following the suggestion from Thompson et al. ^21^ in the TYK2 study), and use a batch size of 30 compounds for exploitation. The ‘random-exploit’ protocol immediately switches to exploitation from the first cycle on. Both the ‘random-explore-exploit’ and ‘random-random-exploit’ protocols acquire 60 more compounds after the initial selection with the purpose of improving coverage of the chemical space, resulting in an effective initial batch size of 120. More precisely, the ‘random-random-exploit’ protocol selects the additional 60 compounds at random, whereas the ‘random-explore-exploit’ protocol utilizes the model’s prediction uncertainty to select the additional compounds over two exploration cycles. Due to the constraint on the total number of acquired compounds, the ‘random-explore-exploit’ and ‘random-random-exploit’ protocols acquire fewer compounds over the exploitation phase than ‘random-exploit’.

**Figure 4:**
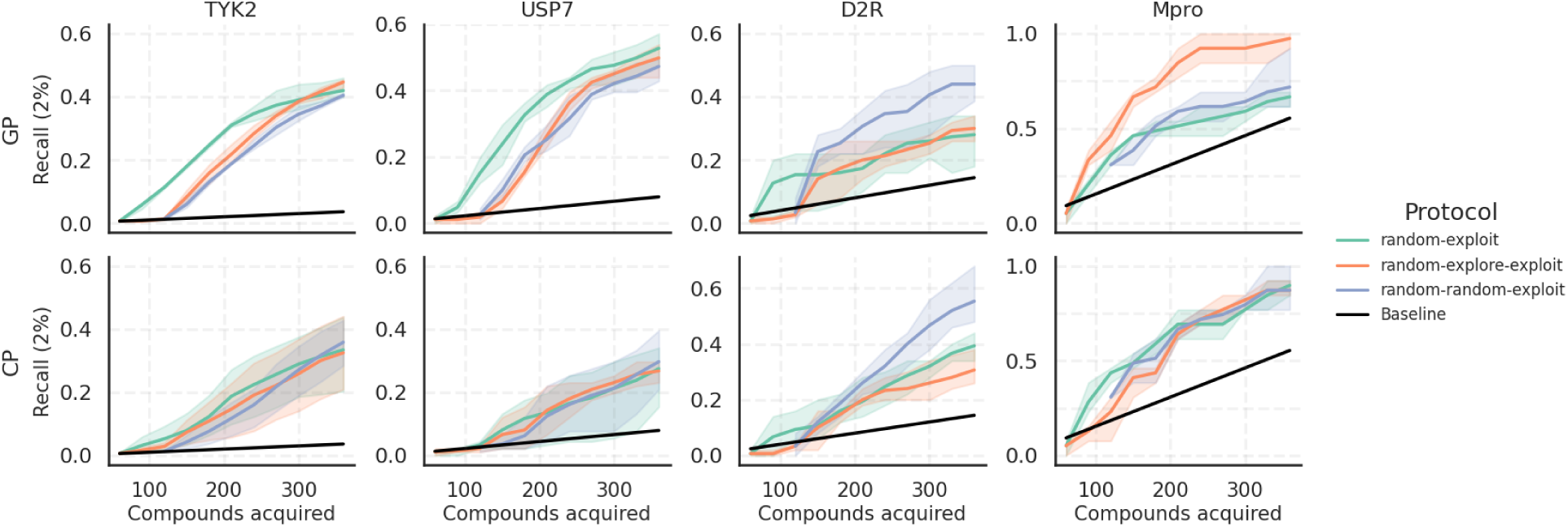
Top 2% Recall achieved with different AL protocols for initial sample selection across the four target datasets. The ‘random-exploit’ protocol acquires 60 compounds at random before switching to exploitation, ‘random-explore-exploit’ acquires another 60 compounds using the prediction uncertainty following the initial batch selected at random, and ‘random-random-exploit’ starts with 120 compounds selected at random. The shaded area displays the variance over three repeats initialized with a different random seed for the initial sample selection. The baseline shows the expectation value for the Recall upon random acquisition. The protocol yielding the best Recall is consistent between the two models for each dataset, but not between different datasets.

Fig. 4 shows the top 2% Recall (Eq. 2) across the four datasets for the three protocols and both GP and CP models, and the top 5% Recall is shown in Fig. S2. Fig. 5A shows the corresponding Spearman *ρ* of the final models trained on all 360 selected compounds. A more detailed plot showing the Spearman *ρ* as a function of the number of compounds acquired can be found in Fig. S3, and the equivalent plots for the R2 and the RMSE are also shown in Fig. S4 and Fig. S5, respectively. A general observation is that a larger initial batch size of 120 with the ‘random-explore-exploit’ and ‘random-random-exploit’ protocols yields more predictive models and therefore augments the likelihood of pinpointing active compounds, as evidenced by the higher Spearman *ρ* (Fig. 5A) and larger slopes in the Recall curves (Fig. 4 and Fig. S2). The same trend is also seen for R2 and RMSE in Fig. S4 and Fig. S5, respectively. However, the increase in initial batch size comes at the expense of an increased initial training cost for the model. The F1 score (Fig. S6 and Fig. S7) decreases in later cycles for all datasets except TYK2 in response to diminishing precision, which is indicative of an increased cost per identified top compound. This decline is more pronounced for smaller datasets.

**Figure 5:**
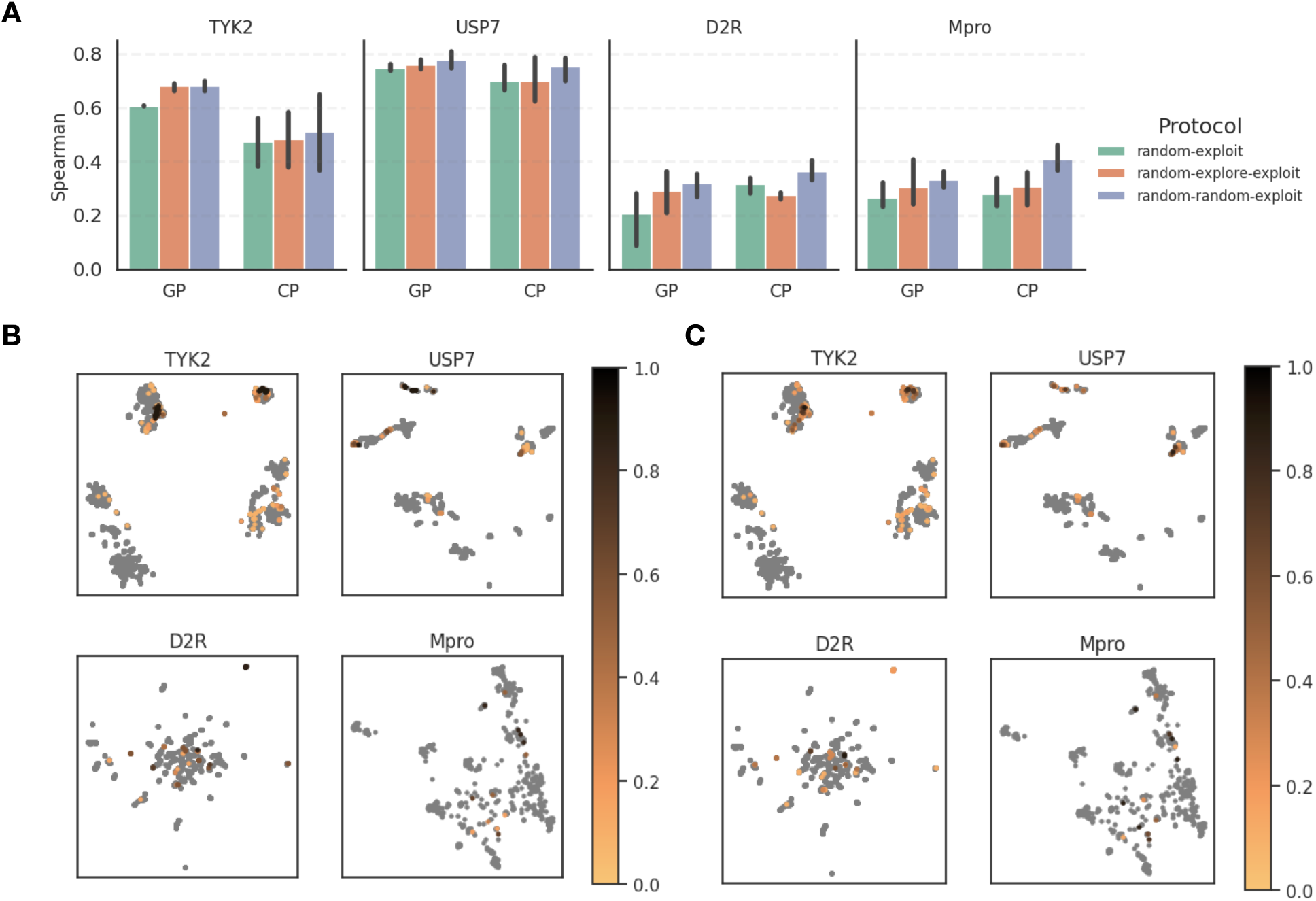
Evaluation of model performance with Spearman *ρ* and UMAP’s showing compound acquisition across AL protocols. **A:** Spearman *ρ* of the final models trained on all 360 selected compounds, emphasizing the enhanced predictability with larger initial batch sizes of 120 in the ‘random-explore-exploit’ and ‘random-random-exploit’ protocols. **B:** UMAP projection of the GP model selection displaying the top 2% compounds in each dataset, colored by the frequency of their acquisition. UMAP shows consistent acquisition of compounds in dense clusters and the challenge of identifying sparse clusters. **C:** UMAP of the CP model, highlighting similar trends in compound acquisition. Overall, we can see that the GP model is more consistent in compound selection than the CP model across multiple AL runs.

For the TYK2 and USP7 datasets, a smaller initial batch size was found to be more adequate with GP, given that the explorative protocols only caught up after around 300 acquired compounds. In contrast, the more heterogeneous D2R and Mpro datasets greatly benefited from a larger initial batch. As can be seen from Fig. S1, D2R predominantly contains top compounds in small dense clusters, while Mpro’s top compounds are more dispersed as singletons. This distinction can explain the opposite trends in terms of selection protocols for these two datasets. However, such nuances are not known *a priori*and cannot be used to inform the choice of an exploration method. The drop of predictive power (Fig. S2 and Fig. S4) on Mpro over the course of AL is associated with the small size of this dataset compared to the number of acquired training samples as well as the exploitative acquisition, which causes a potency imbalance between training and pool data. In comparison to the GP, CP underperformed on all datasets except D2R, although its predictive performance tends to be more consistent between the three protocols. By contrast, CP outperformed GP on D2R, where the underlying global model is likely to account well for the diversity of these compounds.

Fig. 5B and C show UMAPs where the top 2% compounds in each dataset are colored by the frequency of their acquisition (averaged over different protocols and random seeds for initialization) using GP or CP, respectively. A breakdown by protocol for each dataset can also be found in Fig. S8-11, which shows that there is little difference between compounds identified by different protocols. A common trend in Fig. 5B and Fig. 5C is a consistent acquisition of compounds in dense clusters, while sparse clusters remain difficult to track with both models. Overall, we could see that the selection with the GP model is slightly more consistent across multiple AL runs.

We also evaluated the performance of AL relative to training models on a larger proportion of the data, as was the case in the model benchmarking section. The recall achieved by AL using a smaller fraction of the data (Fig. 4) is comparable or better than the models trained in a single batch on 20% of each dataset (Fig. 3), demonstrating the benefits of partitioning the training process into multiple stages. On the other hand, R2 and Spearman *ρ* are lower for models constructed using AL (except for the Mpro dataset due to its size). It might be tempting to conclude that the size of the training set is the deciding factor for a model’s predictive quality, but this is not entirely true. Fig. S3 and Fig. S4 show that the R2 and Spearman *ρ* stop improving after the start of the exploitation phase, and the effective performance of the final models is comparable to the models constructed only from the initial 60 or 120 samples, respectively. Similarly, the models constructed from a large randomly chosen batch are better to account for the variety in the data than samples selected using an exploitative strategy. As stated before, the amount of diversity in the data is strongly linked to the speed of convergence of a model with an increasing number of training samples. On the TYK2 and USP7 datasets, GP models trained on 360 samples reach more than 70% of the rank ordering performance of corresponding models trained on more than twice as many samples (CP underperforms on TYK2, reaching only about 50% of the performance of the larger model). By contrast, this ratio drops to 30-50% for D2R, which also benefited greatly from an increase in the initial batch size. In summary, AL yields greater benefits for compound pools which display a high degree of similarity and strong structure-activity relationships.

#### Batch size for exploitation

To systematically investigate the influence of the batch size in exploitation cycles, we fixed the initial sample selection at 60 samples chosen randomly and the subsequent 60 samples selected based on the exploration strategy. We then evaluated four distinct batch sizes for the subsequent AL cycles, namely 20, 30, 60, and 120. Given our constraint of acquiring a total of 360 compounds, smaller batch sizes necessitate a greater number of AL cycles to reach this total. The protocols varied in batch sizes, with “batch-size-20” using three exploration batches of 20 each followed by twelve exploitation batches of 20, “batch-size-30” using two exploration and eight exploitation batches of 30, “batch-size-60” using one exploration and four exploitation batches of 60, and “batch-size-120” combining one exploration batch of 60 with two exploitation batches of 120.

**Figure 6:**
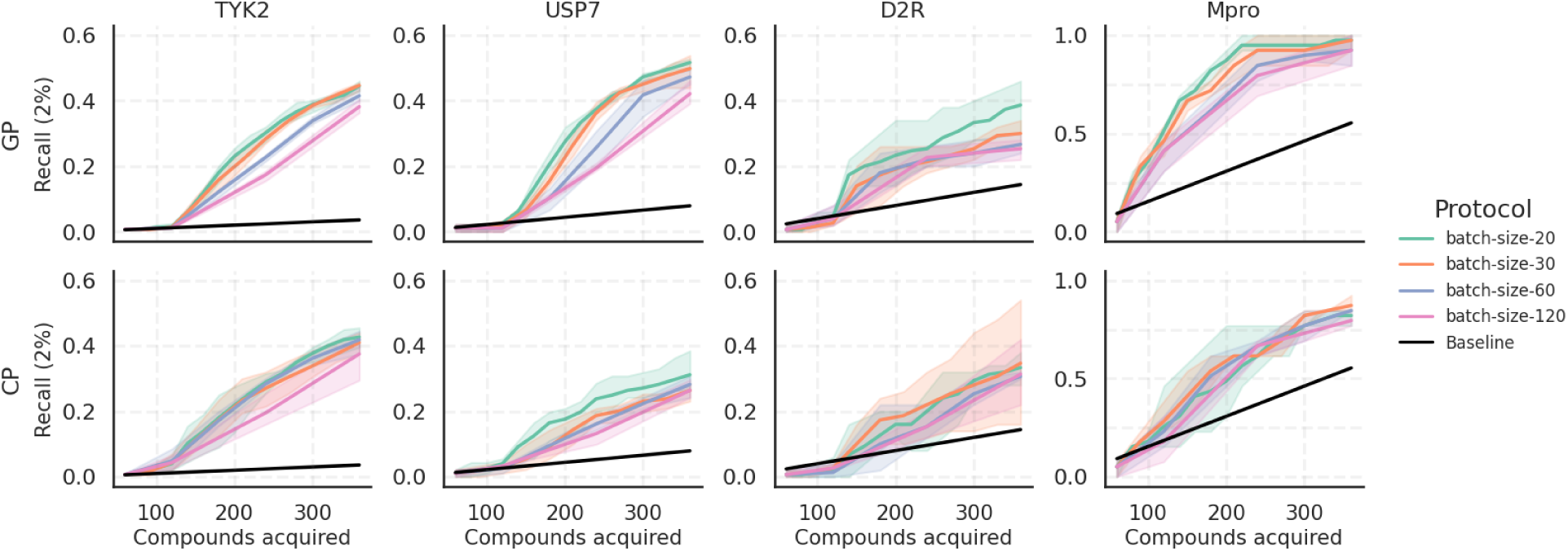
Comparison of the top 2% Recall across AL protocols varying in their batch size. The “batch-size-20” protocol uses three exploration batches and twelve exploitation batches of 20; “batch-size-30” employs two exploration and eight exploitation batches of 30; “batch-size-60” employs a single exploration and four exploitation batches of 60; “batch-size-120” employs one exploration batch of 60 and two exploitation batches of 120. The comparison suggests that the Recall improves with decreasing batch size across all datasets with both models.

Fig. 6 showcases the top 2% Recall across different batch sizes. We can see a clear trend where smaller batch sizes consistently outperform larger ones across all datasets, and this holds true for both GP and CP models. The same trend can be seen in Fig. S12, which shows the top 5% Recall, and also considering F1 scores (Fig. S13 and S14), indicating that smaller batch sizes favor both precision and recall. The same trends are reflected by the metrics Spearman *ρ*, R2 and RMSE across all datasets, except for Mpro (as seen in Fig. S15, S16 and S17). This trend can be understood by considering the ratio of the pool and training sizes. When the pool of available data significantly outnumbers the training set, the model benefits from small, incremental additions to its training data, making a batch size of 1 theoretically optimal. This is because each new data point refines the models understanding. However, in practical scenarios, using extremely small batch sizes might not be feasible, as it would require acquiring potency data in a time-consuming serial manner rather than a more efficient parallel approach. This advantage of smaller batches diminishes when the pool size is roughly equal to the training size, where the predictive power of a model decreases upon imbalancing the data by using an exploitative acquisition strategy.

### Modeling noise on labels

The measurement of binding affinities by experimental and computational means is subject to noise, although being fed to the model as “true” values. To systematically study the influence of noise in training data, we introduced Gaussian noise to each dataset. The noise was generated by sampling random numbers from a Gaussian distribution with a mean of zero and a standard deviation (*σ*) ranging from zero up to two times the standard deviation of the underlying potency data, meaning that the amounts of noise vary between datasets in absolute terms. Our investigation considers the noise multipliers 0, 0.5, 1, 1.5, and 2. Similar to the batch size modulation, we fixed the initial sample selection at 60 samples chosen randomly and the subsequent 60 samples selected using the exploration strategy. We ran the AL pipeline three times with different random seeds for the selection of the initial batch. The transformation of the potency distribution at each noise level for the TYK2 dataset is visually represented in Fig. 7A. Similar transformations for other datasets, namely USP7, D2R, and Mpro, are shown in Fig. S18A, Fig. S19A, and Fig. S20A, respectively, and the UMAPs of the noisy data are shown in Fig. S21. An overarching observation from Fig. S21 is that the introduction of stochastic noise retains the macroscopic structure of the clusters while smoothing out the microscopic features within these clusters across all target datasets.

**Figure 7:**
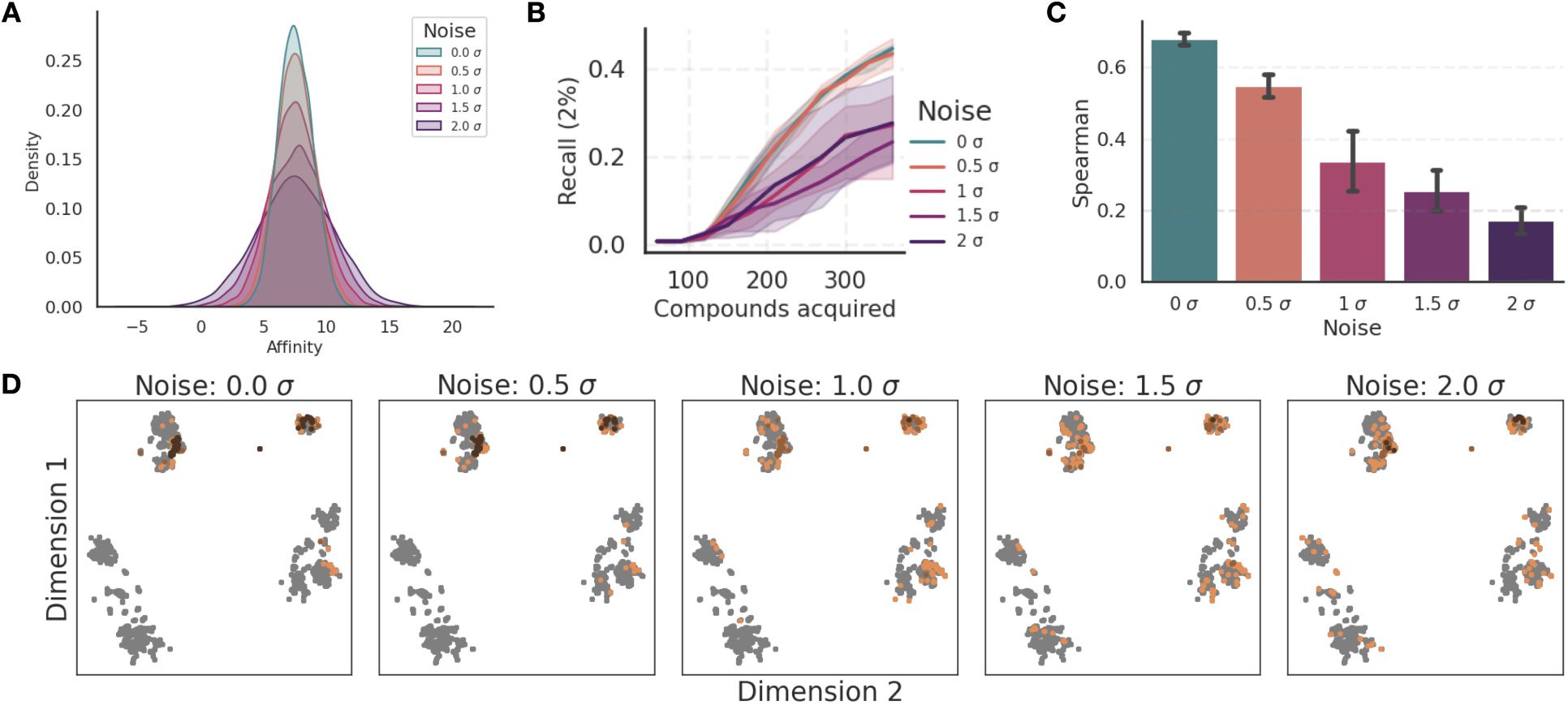
Analysis of the influence of Gaussian noise on the outcomes of AL using the GP model on the TYK2 dataset. The standard deviation of the added Gaussian noise was scaled with respect to the standard deviation of TYK2 pK*_i_* values, with factors ranging from 0 (no noise) to 2. **A:** Kernel Density Estimation plot of the pK*_i_* distribution across varying noise magnitudes. **B:** Top 2% Recall, highlighting a noticeable decline at increased noise levels. **C:** Spearman *ρ* revealing diminished model predictability with increasing noise. **D:** UMAP visualization of the compounds selected in the exploitation phase, colored by the acquisition frequency across three distinct AL iterations with randomized initializations. The UMAPs emphasize the AL framework’s capability to consistently identify top-binding compound clusters, even amidst noise interference.

The introduction of noise has a pronounced effect on the regression performance and Recall of the constructed model. As depicted in Fig. 7B, the top 2% Recall rapidly decays when the noise level exceeds 1 *σ*. This trend of declining Recall with increasing noise is consistently observed across other datasets, as seen in Fig. S18B for USP7, Fig. S19B for D2R, and Fig. S20B for Mpro.

Furthermore, Spearman *ρ* drops with increasing noise levels, as shown in Fig. 7C. However, the models remain predictive even with high amounts of noise, maintaining a positive Spearman *ρ* coefficient. This trend also holds true for USP7 and D2R (Fig. S18C and Fig. S19C), but not for Mpro, where Spearman *ρ* drops below zero for a noise addition of 2 *σ*, as seen in Fig. S20C.

AL can robustly identify relevant clusters even under noisy conditions. Fig. 7D shows compounds acquired in the exploitation phase, colored by the fraction of three AL repeats that selected them. Similar observations can be made for other datasets, as seen in Fig. S18D for USP7, Fig. S19D for D2R, and Fig. S20D for Mpro. These findings are consistent with the work of Bellamy et. al. ^38^ for synthetic affinity data, who found a better Recall in terms of the true (noiseless) labels than considering the noisy values used to train the model. This robustness underscores the fact that while noise may introduce perturbations, learning the overarching structure-activity relationship trends remain preserved. Interestingly, the presence of noise might even be beneficial for exploration, aiding in overcoming potency barriers in the chemical space. However, while noise can enhance exploration, it hampers the exploitative power of the model.

A similar trend of decaying Recall with increasing noise also is observed for the CP model, as depicted in Fig. S22. Spearman *ρ* for the CP model also exhibits a decline with increasing noise levels ( Fig. S23). However, the CP model is far less consistent in identifying relevant clusters compared to GP in the presence of noise (Fig. S24), especially for TYK2 and USP7. In conclusion, while the presence of stochastic noise impairs the performance of AL, the models demonstrate a commendable ability to identify large-scale trends. However, the Gaussian noise simulation in this study primarily accounts for stochastic noise. Systematic errors are not accounted for. They can particularly be detrimental as they introduce a bias in the activity landscape, potentially misguiding the AL process.

## Conclusion and outlook

We comprehensively evaluated the effect of different parameters for AL protocols based on a range of metrics. These capture the identification of top binders, the predictive quality of the underlying ML model, as well as a qualitative analysis of the identified clusters in chemical space. Simulating identical AL runs on different binding affinity datasets allowed us to assess different aspects of the AL strategies, as well as capture trends with respect to the size and composition of a dataset.

Using RBFE to label a chemical library of 5000 compounds, each calculation averaging 8 GPU hours at a rate of $2.50 per GPU hour, results in a total cost of $100K. Using AL and RBFE to label just 300 compounds incurs a cost of $6k and can yield a comparably predictive model. AL is able to identify top binders using a significantly smaller fraction of the data to train. We observed a pronounced dependence on the dataset size, although the diversity of compounds in a dataset is the most decisive factor contributing to the margin of profit achievable by AL. A similar trend was identified when varying the initial batch size for AL, where the more diverse D2R and Mpro datasets benefited from a large initial exploration phase compared to TYK2 and USP7.

Our results suggest that certain design strategies will help in designing the most useful AL protocols. These findings contain implications for the design of ligand pools for active learning. The TYK2 and USP7 datasets, which consistently led to predictive models and a high Recall, consist of few distinct scaffolds and a large number of substitutions, or even a combinatorial enumeration of substituents in the case of TYK2. The confinement of chemical space is essential for a small number of samples to be able to represent a sizable pool. In practice, however, a combinatorial exploration of substituents is not always desirable when a strict filtering of compounds is necessary due to constraints on the physicochemical properties. In such cases, increasing the batch size is a more defensive approach to achieve success with AL. In the present study, using the model uncertainty in the initial exploration phase did not yield any observable benefits over selecting the compounds at random.

For the exploitation phase, our findings consistently suggest that training in small batches results in the highest Recall. The performance gains, however, are incremental when reducing the batch size below 30 samples. Very small batches are also undesirable from a practical perspective, due to the increase in the number of AL cycles and the overall turnaround time. The addition of noise to potency data was found to have detrimental effects on the exploitative power of AL if the variance of the noise is equal to or larger than the variance of the affinity values. On the other hand, the noise did not prevent the GP model from finding large-scale active regions in the chemical space, even when its variance exceeded the underlying signal. The CP model was more affected by noise in the data and lost its predictive power upon high levels of noise. Together with the fact that CP outperformed GP on the AL runs for D2R, whereas GP performed better in the general case, it is likely to be more sensitive to the local structure of chemical space and at the same time, is more vulnerable to noise.

Using a Gaussian model noise is a drastic simplification to account for the entirety of RBFE errors, and understanding their nature for a given ligand series is crucial. A validation prior to running AL-RBFE is highly recommended, as it may not only provide insight into the magnitude of stochastic errors relative to the dynamic range of RBFE values but also reveal systematic offsets for certain functional groups, which can alter the entire course of an AL run.

Understanding the relationships between data, models, and selection strategies in AL pipelines paves the way to establishing protocols for choosing these parameters in an automated fashion. Using the findings from this work to parameterize AL pipelines based on distances between data points and the retrospective evaluation of the models performance is a promising avenue for future work.

## Data Availability

All data for the experiments carried out can be found at https://github.com/meyresearch/ActiveLearning_BindingAffinity.

## Acknowledgement

This work was supported by the United Kingdom Research and Innovation (grant EP/S02431X/1), UKRI Centre for Doctoral Training in Biomedical AI at the University of Edinburgh, School of Informatics and Exscientia Plc, Oxford.

## References

1. Reker, D. Practical considerations for active machine learning in drug discovery. Drug Discov. Today 2019, 32–33, 73–79.

2. Graff, D. E.; Shakhnovich, E. I.; Coley, C. W. Accelerating high-throughput virtual screening through molecular pool-based active learning. Drug Discov. Today 2015, 20, 458–465.

3. Yu, J.; Li, X.; Zheng, M. Current status of active learning for drug discovery. Artif. Intell. Life Sci. 2021, 1, 100023.

4. Ahmadi, M.; Vogt, M.; Iyer, P.; Bajorath, J.; Fröhlich, H. Predicting Potent Compounds via Model-Based Global Optimization. J. Chem. Inf. Model. 2013, 53, 553–559.

5. Varela, R.; Walters, W. P.; Goldman, B. B.; Jain, A. N. Iterative Refinement of a Binding Pocket Model: Active Computational Steering of Lead Optimization. J. Med. Chem. 2012, 55, 8926–8942.

6. Fusani, L.; Cabrera, A. C. Active learning strategies with COMBINE analysis: new tricks for an old dog. J. Comput. Aided Mol. Des. 2019, 33, 287–294.

7. Desai, B.; Dixon, K.; Farrant, E.; Feng, Q.; Gibson, K. R.; Hoorn, W. P. v.; Mills, J.; Morgan, T.; Parry, D. M.; Ramjee, M. K.; Selway, C. N.; Tarver, G. J.; Whitlock, G.; Wright, A. G. Rapid Discovery of a Novel Series of Abl Kinase Inhibitors by Application of an Integrated Microfluidic Synthesis and Screening Platform. J. Med. Chem. 2013, 56, 3033–3047.

8. Stanzione, F.; Giangreco, I.; Cole, J. C. Use of molecular docking computational tools in drug discovery. Prog. Med. Chem. 2021, 60, 273–343.

9. Wang, R.; Lu, Y.; Wang, S. Comparative evaluation of 11 scoring functions for molecular docking. J. Med. Chem. 2003, 46, 2287–2303.

10. Mey, A. S.; Allen, B. K.; Macdonald, H. E. B.; Chodera, J. D.; Hahn, D. F.; Kuhn, M.; Michel, J.; Mobley, D. L.; Naden, L. N.; Prasad, S.; Rizzi, A.; Scheen, J.; Shirts, M. R.; Tresadern, G.; Xu, H. Best Practices for Alchemical Free Energy Calculations. Living J. Mol. Sci. 2020, 2, 18378.

11. Hahn, D. F.; Bayly, C. I.; Boby, M. L.; Macdonald, H. E. B.; Chodera, J. D.; Gapsys, V.; Mey, A. S.; Mobley, D. L.; Benito, L. P.; Schindler, C. E.; Tresadern, G.; Warren, G. L. Best practices for constructing, preparing, and evaluating protein-ligand binding affinity benchmarks. Living J. Mol. Sci. 2022, 4, 1497–1497.

12. Kimber, T. B.; Chen, Y.; Volkamer, A. Deep learning in virtual screening: recent applications and developments. Int. J. Mol. Sci. 2021, 22, 4435.

13. Lyu, J.; Irwin, J. J.; Shoichet, B. K. Modeling the expansion of virtual screening libraries. Nat. Chem. Biol. 2023, 19, 712–718.

14. Gorantla, R.; Kubincova, A.; Weisse, A. Y.; Mey, A. S. J. S. From Proteins to Ligands: Decoding Deep Learning Methods for Binding Affinity Prediction. bioRxiv 2023,

15. Graff, D. E.; Shakhnovich, E. I.; Coley, C. W. Accelerating high-throughput virtual screening through molecular pool-based active learning. Chem. Sci. 2021, 12, 7866– 7881.

16. Fujiwara, Y.; Yamashita, Y.; Osoda, T.; Asogawa, M.; Fukushima, C.; Asao, M.; Shimadzu, H.; Nakao, K.; Shimizu, R. Virtual Screening System for Finding Structurally Diverse Hits by Active Learning. J. Chem. Inf. Model. 2008, 48, 930–940.

17. Yang, Y.; Yao, K.; Repasky, M. P.; Leswing, K.; Abel, R.; Shoichet, B. K.; Jerome, S. V. Efficient Exploration of Chemical Space with Docking and Deep Learning. J. Chem. Theory Comput. 2021, 17, 7106–7119.

18. Berenger, F.; Kumar, A.; Zhang, K. Y. J.; Yamanishi, Y. Lean-Docking: Exploiting Ligands’ Predicted Docking Scores to Accelerate Molecular Docking. J. Chem. Inf. Model. 2021, 61, 2341–2352.

19. Konze, K.; Bos, P. H.; Dahlgren, M. K.; Leswing, K.; Brohman, I. T.; Bortolato, A.; Robbason, B.; Abel, R.; Bhat, S. Reaction-based Enumeration, Active Learning, and Free Energy Calculations to Rapidly Explore Synthetically Tractable Chemical Space and Optimize Potency of Cyclin-dependent Kinase 2 Inhibitors. J. Chem. Inf. Model. 2019, 59, 3782–3793.

20. Ghanakota, P.; Bos, P. H.; Konze, K. D.; Staker, J.; Marques, G.; Marshall, K.; Leswing, K.; Abel, R.; Bhat, S. Combining Cloud-Based Free-Energy Calculations, Synthetically Aware Enumerations, and Goal-Directed Generative Machine Learning for Rapid Large-Scale Chemical Exploration and Optimization. J. Chem. Inf. Model. 2020, 60, 4311–4325.

21. Thompson, J.; Walters, W. P.; Feng, J. A.; Pabon, N. A.; Xu, H.; Goldman, B. B.; Moustakas, D.; Schmidt, M.; York, F. Optimizing active learning for free energy calculations. Artif. Intell. Life Sci. 2022, 2, 100050.

22. Gusev, F.; Gutkin, E.; Kurnikova, M. G.; Isayev, O. Active learning guided drug design lead optimization based on relative binding free energy modeling. J. Chem. Inf. Model. 2023, 63, 583–594.

23. Khalak, Y.; Tresadern, G.; Hahn, D. F.; Groot, B. L. d.; Gapsys, V. Chemical Space Exploration with Active Learning and Alchemical Free Energies. J. Chem. Theory Comput. 2022, 18, 6259–6270.

24. McInnes, L.; Healy, J.; Melville, J. Umap: Uniform manifold approximation and projection for dimension reduction. arXiv:1802.03426 2018,

25. Shen, W.-f.; Tang, H.-w.; Li, J.-b.; Li X.; Chen, S. Multimodal data fusion for supervised learning-based identification of USP7 inhibitors: a systematic comparison. J. Cheminform. 2023, 15, 1–16.

26. Mendez, D. et al. ChEMBL: towards direct deposition of bioassay data. Nucleic Acids Res. 2019, 47, D930–D940.

27. O’Boyle, N. M.; Banck, M.; James, C. A.; Morley, C.; Vandermeersch, T.; Hutchison, G. R. Open Babel: An open chemical toolbox. J. Cheminform. 2011, 3, 1–14.

28. Zhang, Z.; Zhao, B.; Xie, A.; Bian, Y.; Zhou, S. Activity Cliff Prediction: Dataset and Benchmark. 2023; arXiv:2302.07541 (accessed Sep 10, 2023).

29. Achdout, H.; Aimon, A.; Bar-David, E.; Morris, G. COVID moonshot: open science discovery of SARS-CoV-2 main protease inhibitors by combining crowdsourcing, high-throughput experiments, computational simulations, and machine learning. BioRxiv 2020,

30. Schulz, E.; Speekenbrink, M.; Krause, A. A tutorial on Gaussian process regression: Modelling, exploring, and exploiting functions. J. Math. Psychol. 2018, 85, 1–16.

31. Gardner, J.; Pleiss, G.; Weinberger, K. Q.; Bindel, D.; Wilson, A. G. Gpytorch: Black-box matrix-matrix gaussian process inference with gpu acceleration. NeurIPS 2018, 31 .

32. Software, O. S. OEChem TK. 2023; http://www.eyesopen.com, accessed 2023-08-30.

33. Bento, A. P.; Hersey, A.; Félix, E.; Landrum, G.; Gaulton, A.; Atkinson, F.; Bellis, L. J.; De Veij, M.; Leach, A. R. An open source chemical structure curation pipeline using RDKit. J. Cheminform. 2020, 12, 1–16.

34. Yang, K.; Swanson, K.; Jin, W.; Coley, C.; Eiden, P.; Gao, H.; Guzman-Perez, A.; Hopper, T.; Kelley, B.; Mathea, M.; Palmer, A.; Settels, V.; Jaakkola, T.; Jensen, K.; Barzilay, R. Analyzing learned molecular representations for property prediction. J. Chem. Inf. Model. 2019, 59, 3370–3388.

35. Gal, Y.; Ghahramani, Z. Dropout as a bayesian approximation: Representing model uncertainty in deep learning. ICML. 2016; pp 1050–1059.

36. Heid, E.; Greenman, K. P.; Chung, Y.; Li, S.-C.; Graff, D. E.; Vermeire, F. H.; Wu, H.; Green, W. H.; McGill, C. J. Chemprop: A Machine Learning Package for Chemical Property Prediction. 2023,

37. McInnes, L.; Healy, J.; Saul, N.; Grossberger, L. UMAP: Uniform Manifold Approximation and Projection. J. Open Source Softw. 2018, 3, 861.

38. Bellamy, H.; Rehim, A. A.; Orhobor, O. I.; King, R. Batched Bayesian Optimization for Drug Design in Noisy Environments. J. Chem. Inf. Model. 2022, 62, 3970–3981.

