## Supplementary Information for "Benchmarking active learning protocols for ligand binding affinity prediction"


#### Acronyms

- **AL** - Active learning
- **GP** - Gaussian process
- **CP** - Chemprop
- **UMAP** - Uniform Manifold Approximation and Projection
- **RMSE** - Root Mean Square Error

#### Datasets

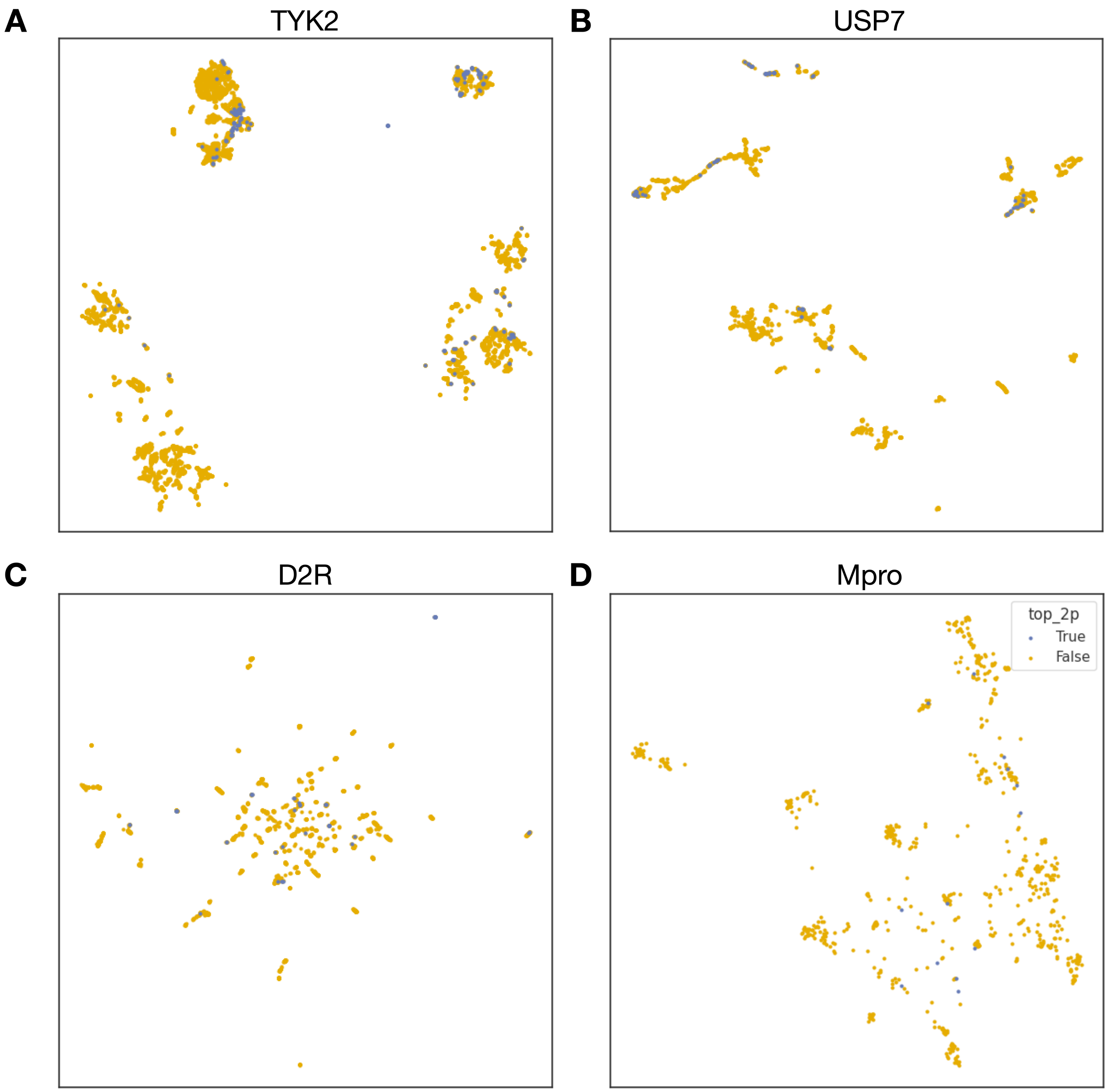


Figure S1: UMAP projection highlighting the top 2% binders in datasets used in our study- **A:** TYK2, **B:** USP7, **C:** D2R, and **D:** Mpro. Blue markers denote the top 2% compounds and the remaining compounds present in the dataset in yellow. We can see that the top 2% compounds for both TYK2 and USP7 targets are present in the dense clusters on the top. In the D2R dataset, the top 2% compounds are present in small clusters, and these small clusters are scattered over the entire chemical space with a few top binders in each. We can see top binders as singletons for Mpro dataset.

### Results of AL strategies

#### Selection of Initial Samples

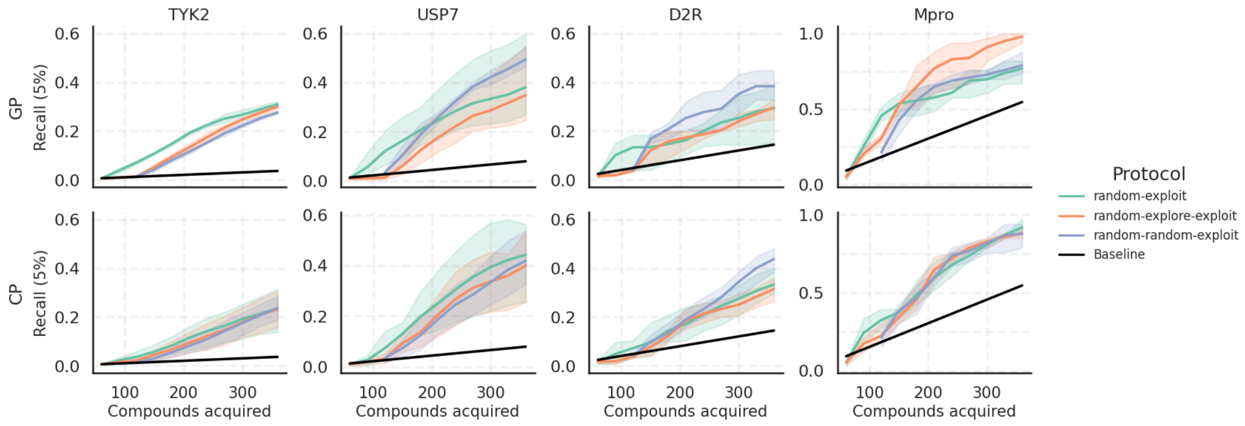


Figure S2: 5% Recall using different AL protocols. Here the compounds acquired shows the number of compounds selected over several AL cycles. The shaded area is variation over 3 AL runs with different seeds, and that baseline is the expectation value given a random selection.

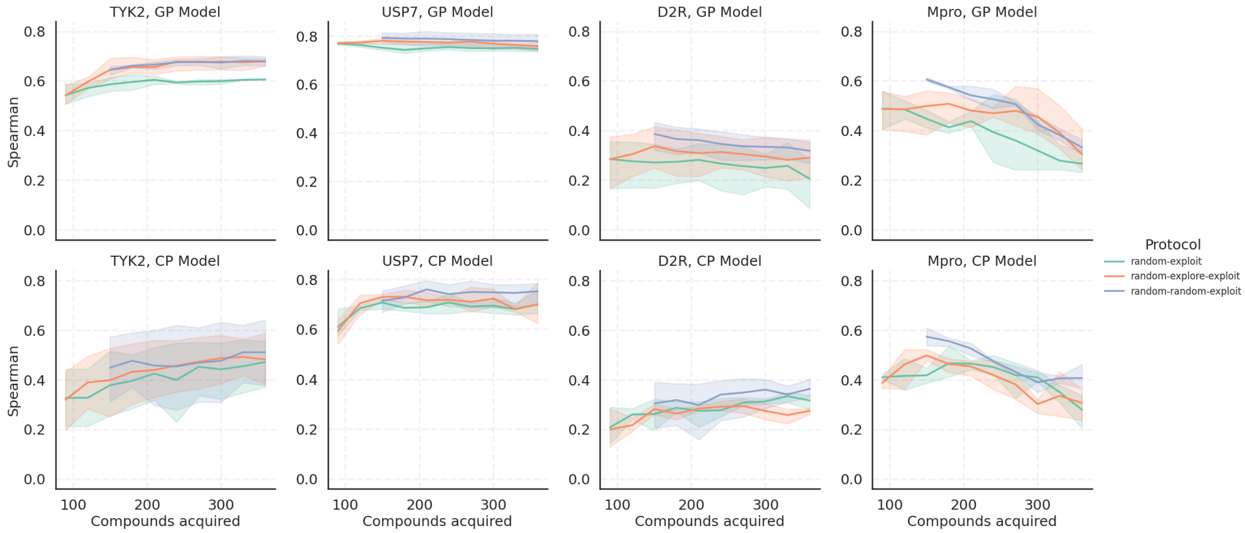


Figure S3: Spearman $\rho$ using different AL protocols on all four target datasets with GP and CP models. Compounds acquired are cumulative over AL cycles. The shaded area is variation over 3 AL runs with different seeds.

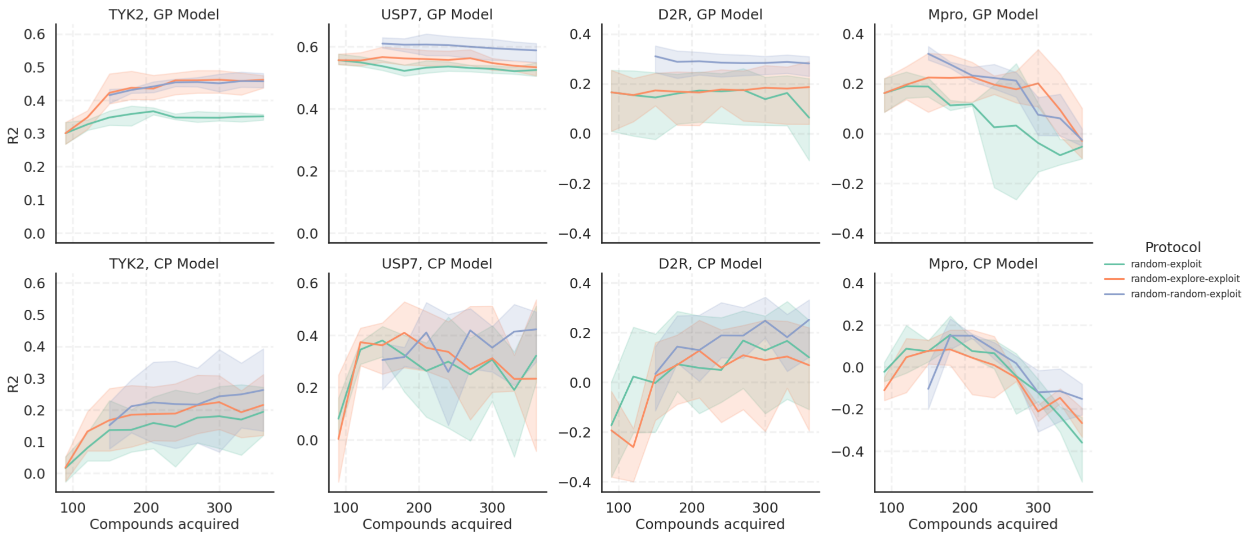


Figure S4: Coefficient of determination, R2 score using different AL protocols on all four target datasets with GP and CP models. Compounds acquired are cumulative over AL cycles. The shaded area is variation over 3 AL runs with different seeds.

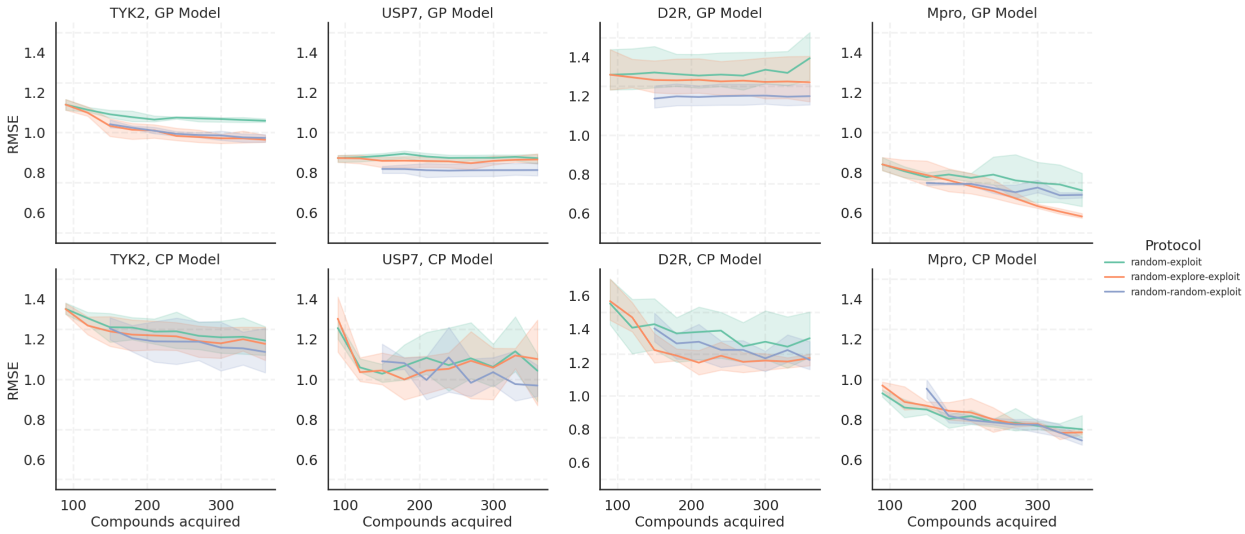


Figure S5: RMSE using different AL protocols on all four target datasets with GP and CP models. Compounds acquired are cumulative over AL cycles. The shaded area is variation over 3 AL runs with different seeds.

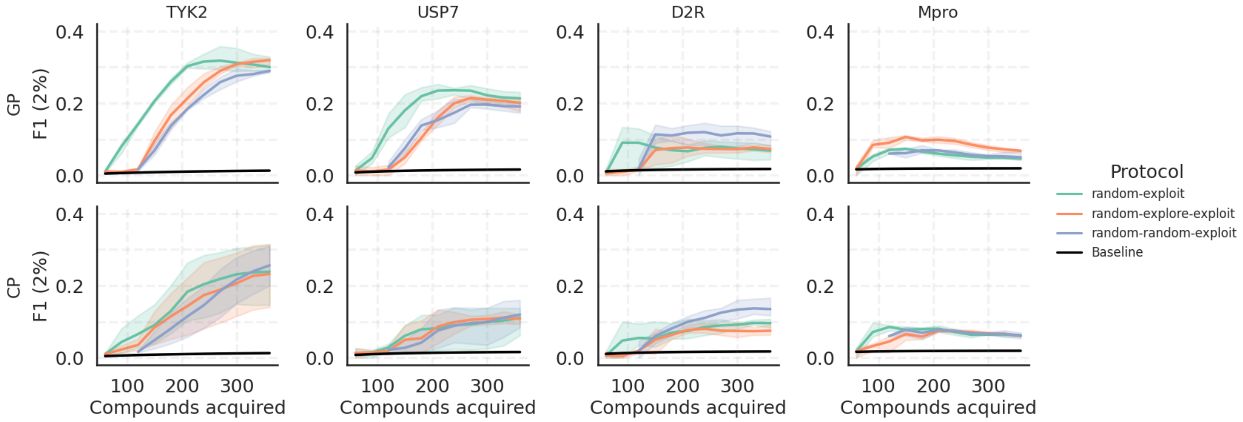


Figure S6: Comparing F1 score for top 2% compounds for GP and CP models using different AL protocols on all four target datasets.

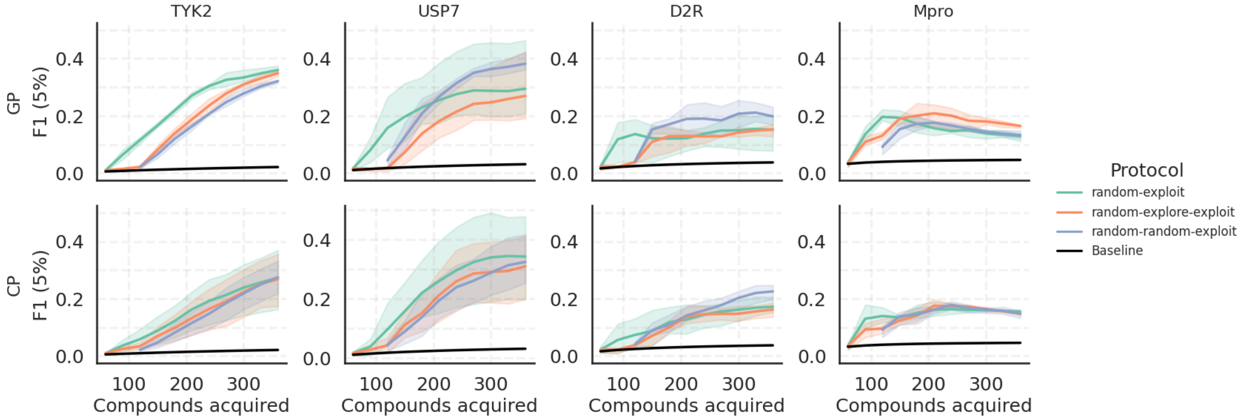


Figure S7: Comparing F1 score for top 5% compounds for GP and CP models using different AL protocols on all four target datasets.

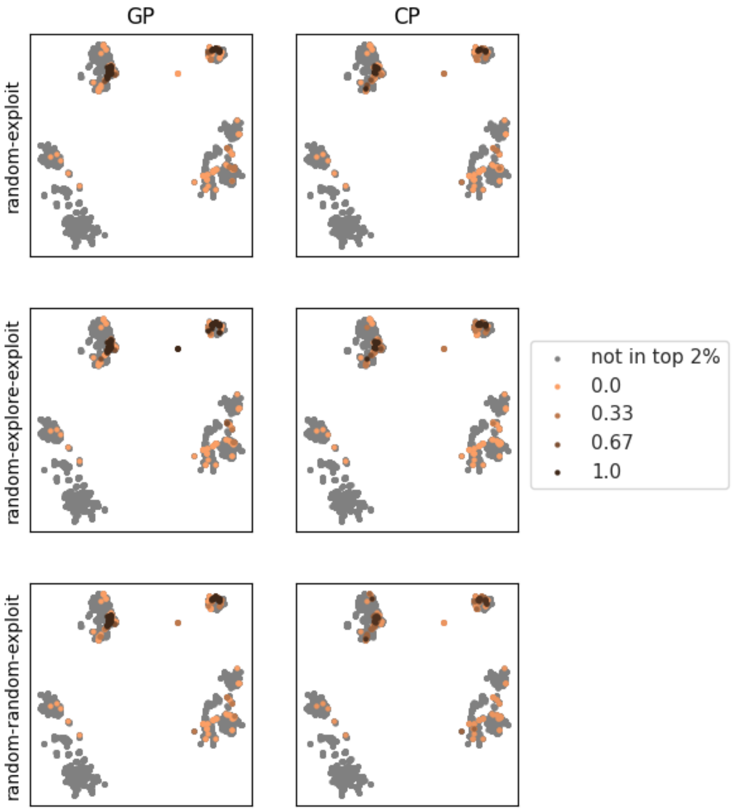


Figure S8: Top 2% TYK2 compounds on UMAP colored by the fraction of the 3 AL runs where they were acquired using GP and CP models.

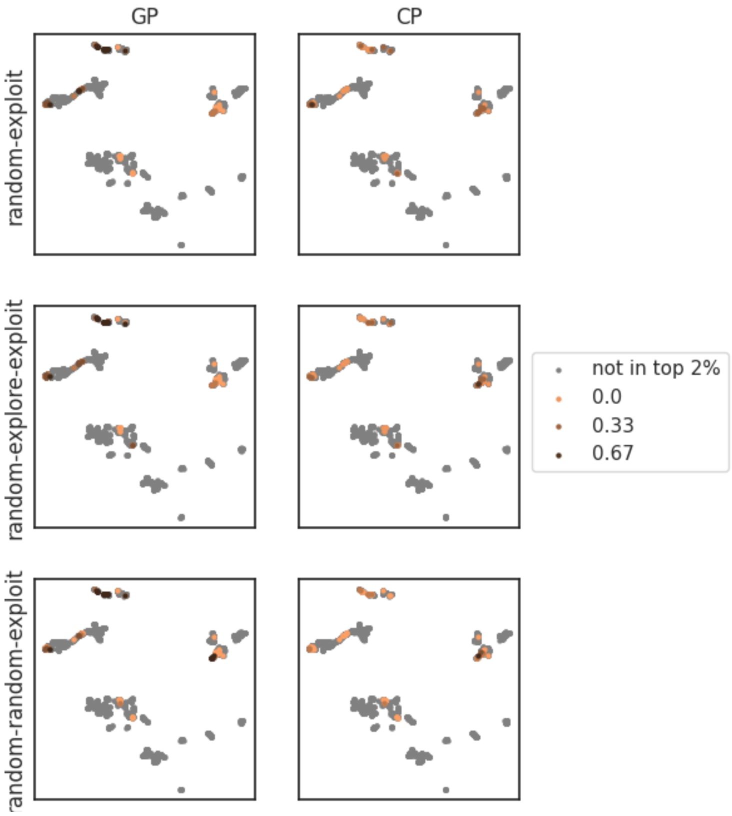


Figure S9: Top 2% USP7 compounds on UMAP colored by the fraction of the 3 AL runs where they were acquired using GP and CP models.

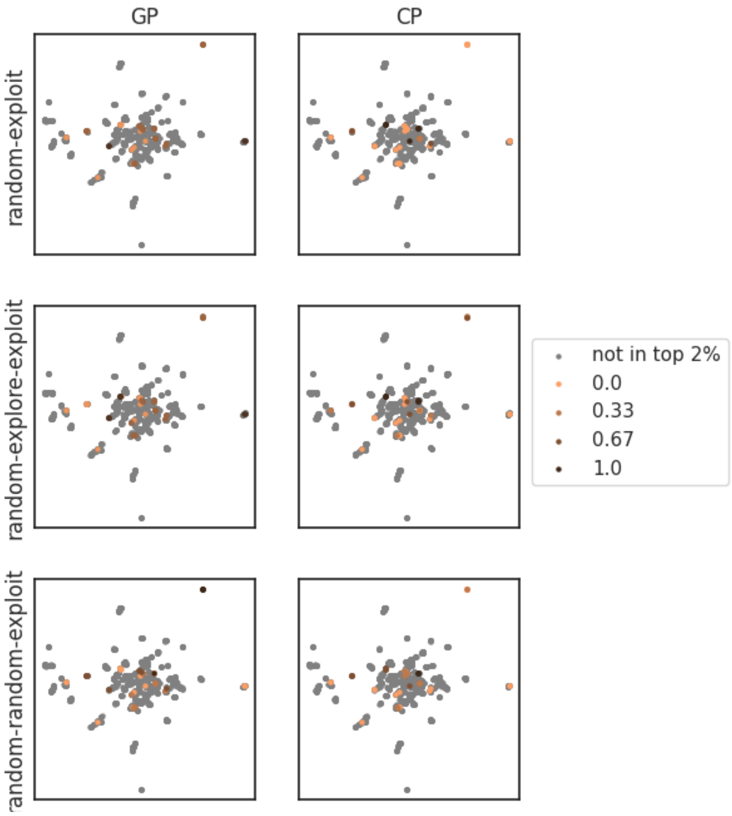


Figure S10: Top 2% D2R compounds on UMAP colored by the fraction of the 3 AL runs where they were acquired using GP and CP models.

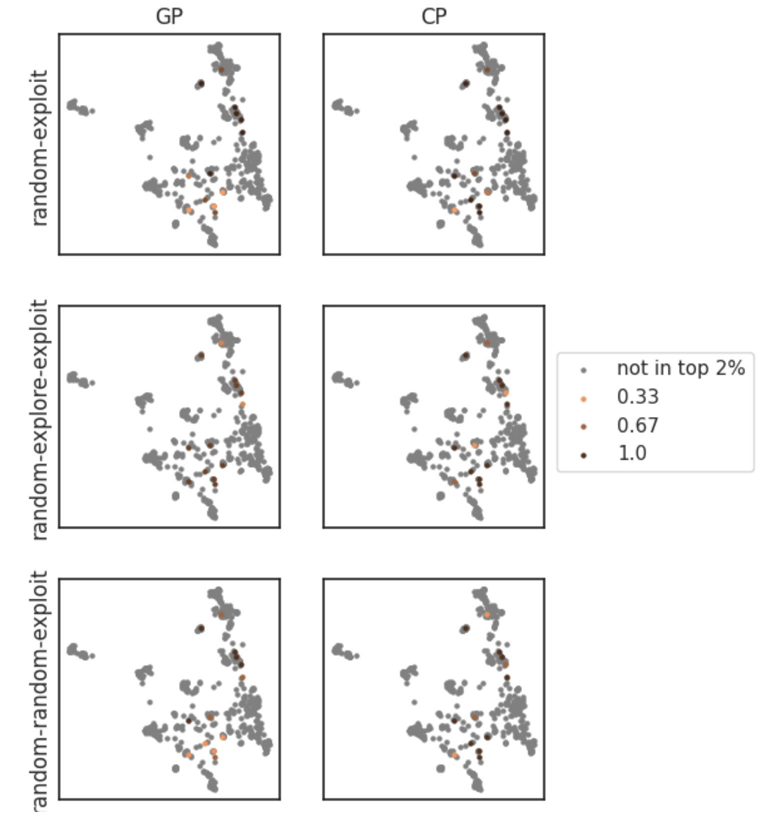


Figure S11: Top 2% Mpro compounds on UMAP colored by the fraction of the 3 AL runs where they were acquired using GP and CP models.

#### Influence of batch size

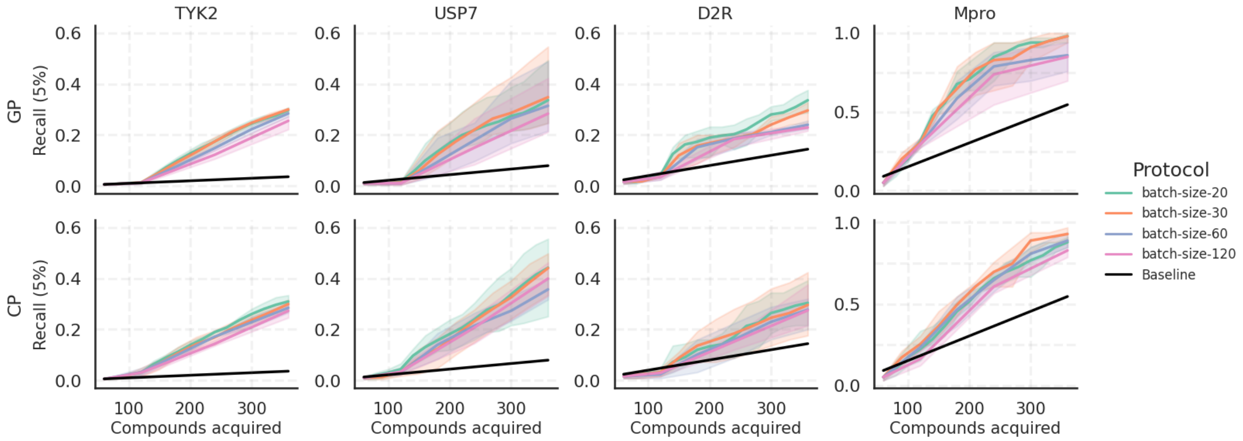


Figure S12: Comparing GP and CP model performance on 5% Recall with different batch sizes. The "batch-size-20" protocol employs three exploration batches and twelve exploitation batches of 20; "batch-size-30" employs two exploration and eight exploitation batches of 30; "batch-size-60" employs a single exploration and four exploitation batches of 60; "batch-size-120" employs one exploration batch of 60 and two exploitation batches of 120. Here the compounds acquired shows the number of compounds selected over several AL cycles. The shaded area is variation over 3 AL runs with different seeds, and that baseline is the expectation value given a random selection.

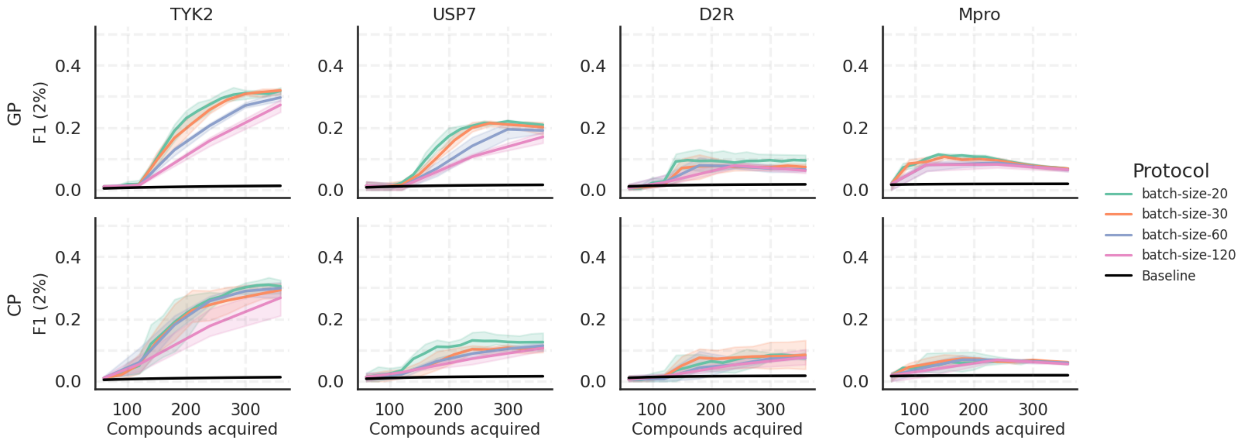


Figure S13: Comparing GP and CP model performance on F1 score (2%) with different batch sizes. Compounds acquired are cumulative over AL cycles. The shaded area is variation over 3 AL runs with different seeds, and that baseline is the expectation value given a random selection.

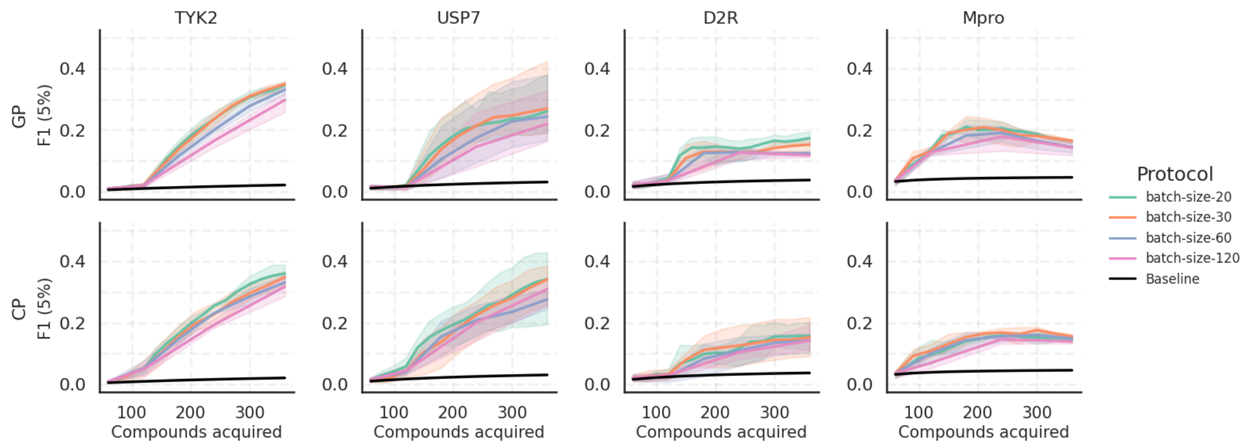


Figure S14: Comparing GP and CP model performance on F1 score (5%) with different batch sizes. Compounds acquired are cumulative over AL cycles. The shaded area is variation over 3 AL runs with different seeds, and that baseline is the expectation value given a random selection.

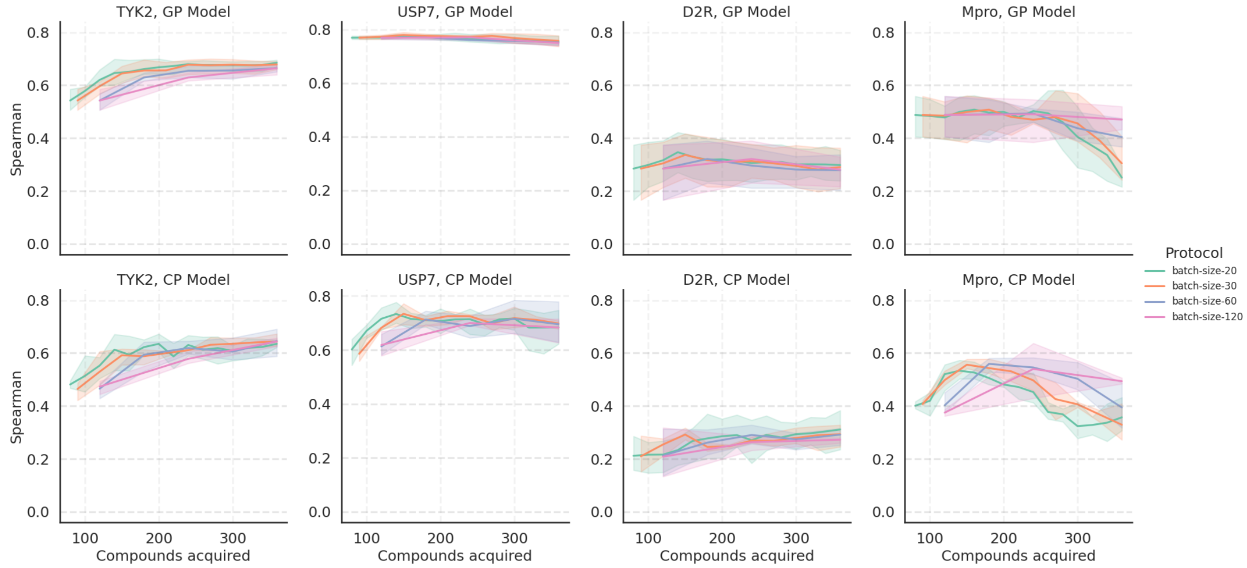


Figure S15: Spearman $\rho$ using different AL protocols with varying batch sizes on all four target datasets with GP and CP models. Compounds acquired are cumulative over AL cycles. The shaded area is variation over 3 AL runs with different seeds.

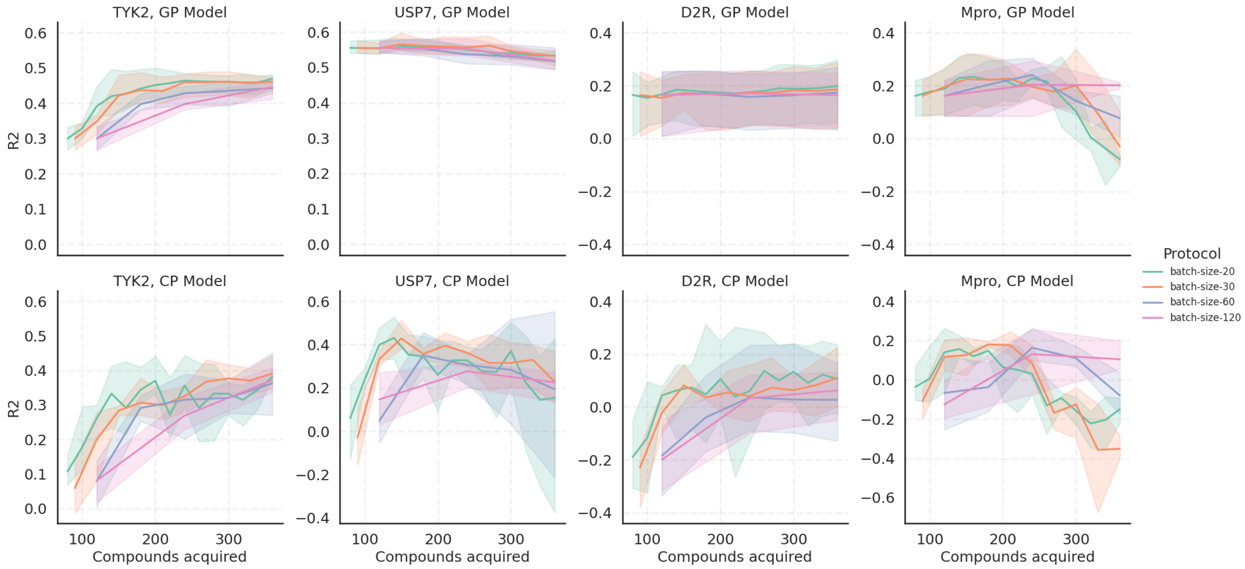


Figure S16: Coefficient of determination, R2 using different AL protocols with varying batch sizes on all four target datasets with GP and CP models. Compounds acquired are cumulative over AL cycles. The shaded area is variation over 3 AL runs with different seeds.

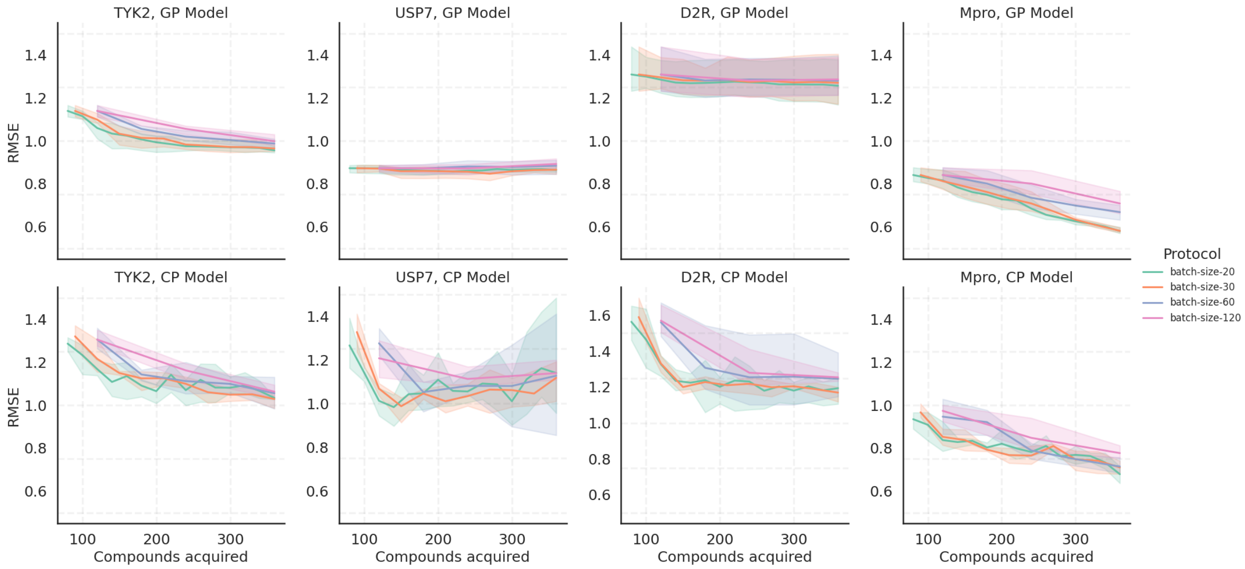


Figure S17: RMSE using different AL protocols with varying batch sizes on all four target datasets with GP and CP models. Compounds acquired are cumulative over AL cycles. The shaded area is variation over 3 AL runs with different seeds.

#### Modelling noise on labels

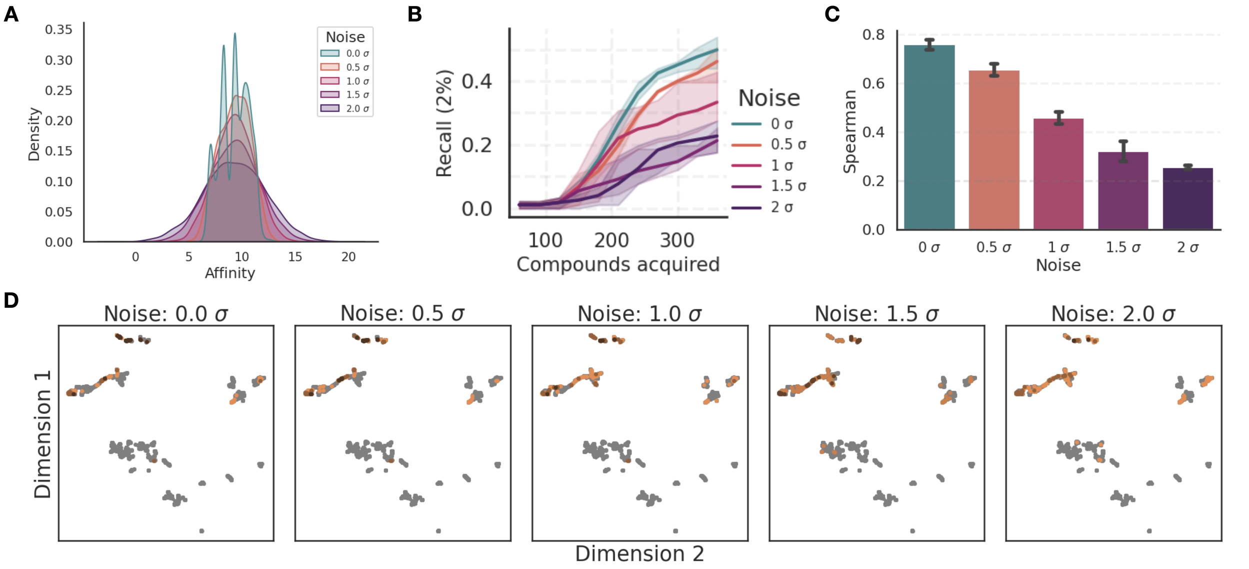


Figure S18: Analysis of the influence of Gaussian noise on the outcomes of AL using the GP model on the USP7 dataset. The standard deviation of the added Gaussian noise was scaled with respect to the standard deviation of USP7 affinities, with factors ranging from 0 (no noise) to 2. **A:** Kernel Density Estimation plot of the affinity score distribution across varying noise magnitudes. **B:** Top 2% Recall, highlighting a noticeable decline at increased noise levels. **C:** Spearman $\rho$ revealing diminished model predictability with increasing noise. **D:** UMAP visualization of the compounds selected in the exploitation phase, colored by the acquisition frequency across three distinct AL iterations with randomized initializations. The UMAPs emphasize the AL framework's capability to consistently identify top-binding compound clusters, even amidst noise interference.

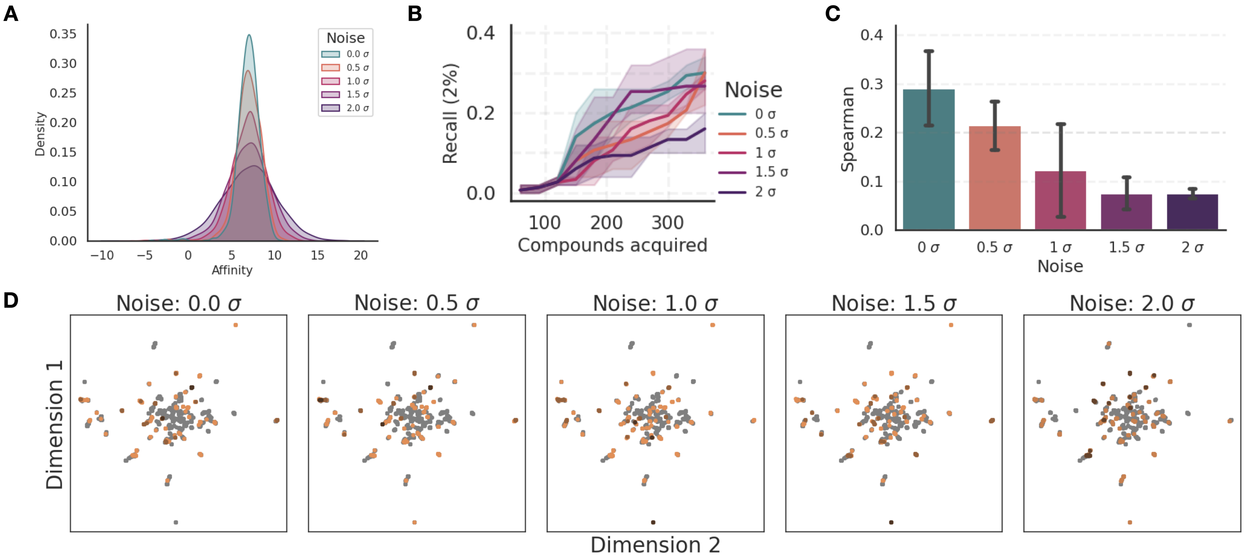


Figure S19: Analysis of the influence of Gaussian noise on the outcomes of AL using the GP model on the D2R dataset. The standard deviation of the added Gaussian noise was scaled with respect to the standard deviation of D2R affinities, with factors ranging from 0 (no noise) to 2. **A:** Kernel Density Estimation plot of the affinity score distribution across varying noise magnitudes. **B:** Top 2% Recall shown at different noise levels. **C:** Spearman $\rho$ shown at different noise levels. **D:** UMAP visualization of the compounds selected in the exploitation phase, colored by the acquisition frequency across three distinct AL iterations with randomized initializations.

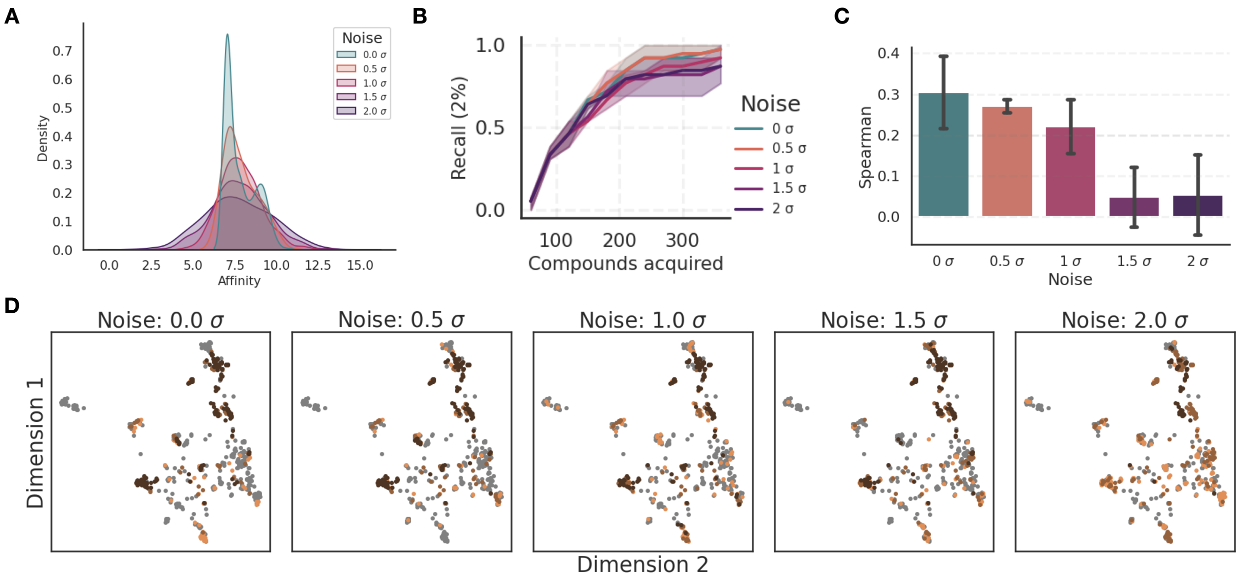


Figure S20: Analysis of the influence of Gaussian noise on the outcomes of AL using the GP model on the Mpro dataset. The standard deviation of the added Gaussian noise was scaled with respect to the standard deviation of Mpro affinities, with factors ranging from 0 (no noise) to 2. **A:** Kernel Density Estimation plot of the affinity score distribution across varying noise magnitudes. **B:** Top 2% Recall shown at different noise levels. **C:** Spearman $\rho$ shown at different noise levels. **D:** UMAP visualization of the compounds selected in the exploitation phase, colored by the acquisition frequency across three distinct AL iterations with randomized initializations.

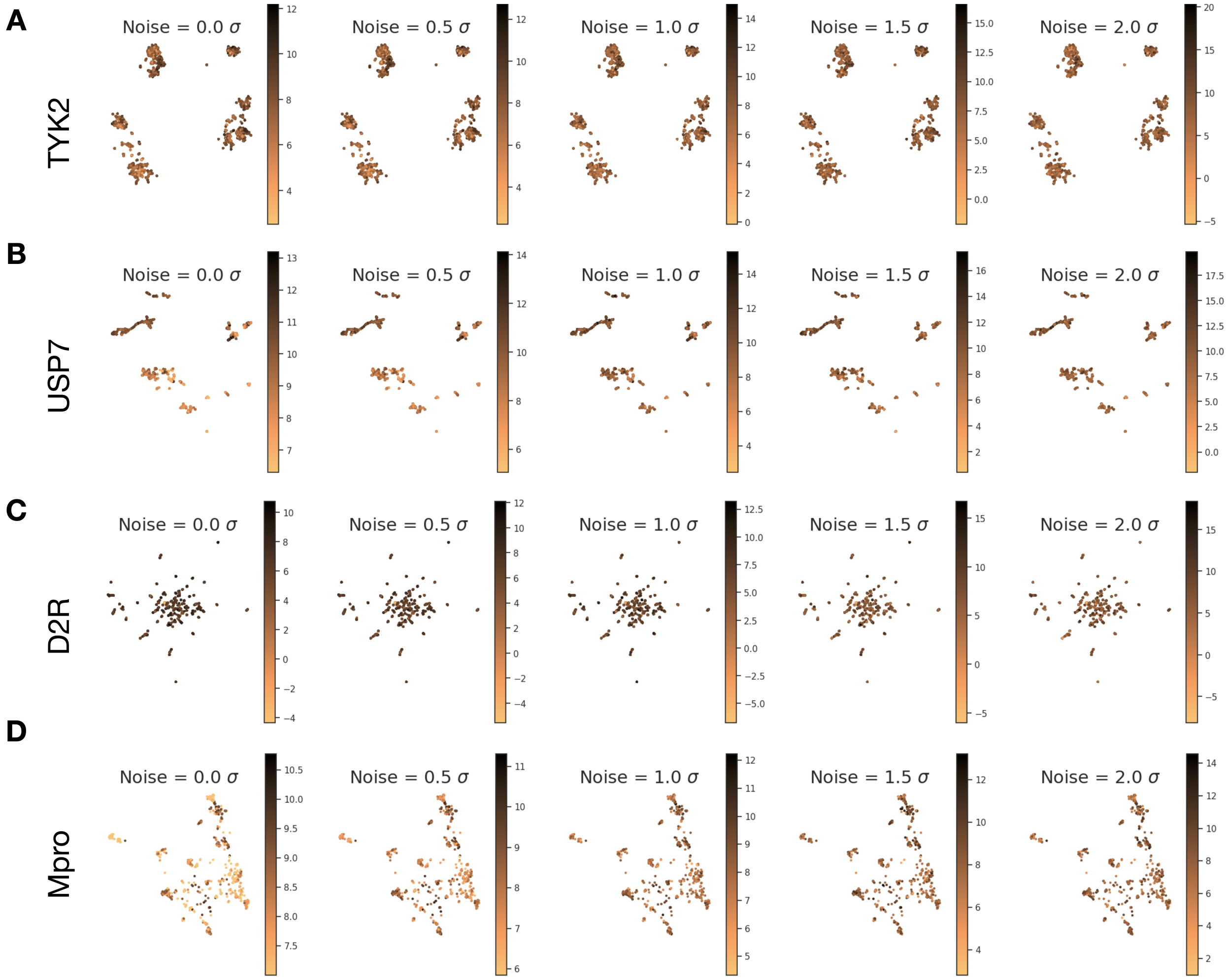


Figure S21: UMAPs showing the chemical space changes at different noise levels on for four datasets used in our study- **A:** TYK2, **B:** USP7, **C:** D2R, and **D:** Mpro. UMAP visualization of the compounds colored by the potency label at different noise levels.

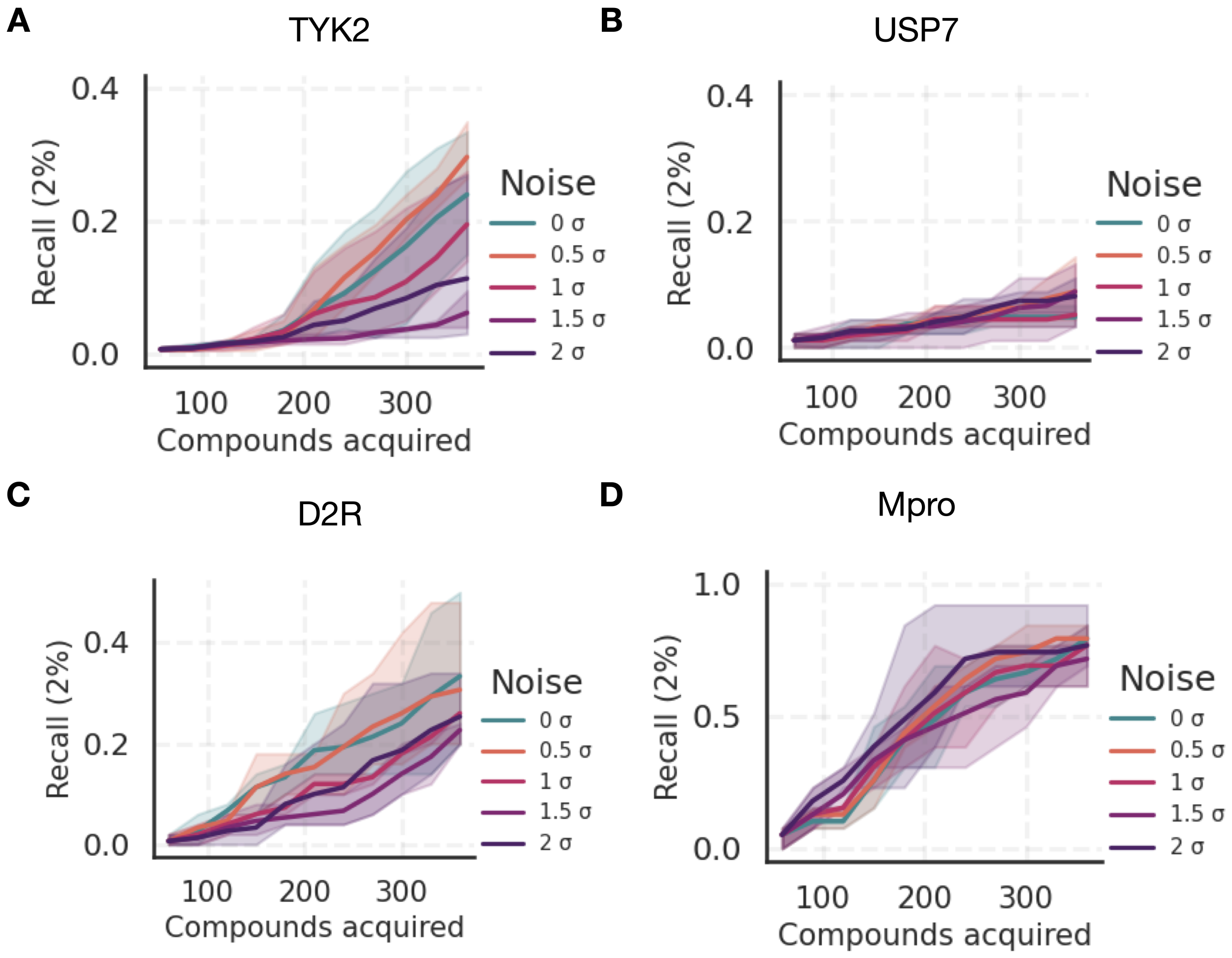


Figure S22: Top 2% Recall with CP models at different noise levels on for four datasets used in our study- **A:** TYK2, **B:** USP7, **C:** D2R, and **D:** Mpro.

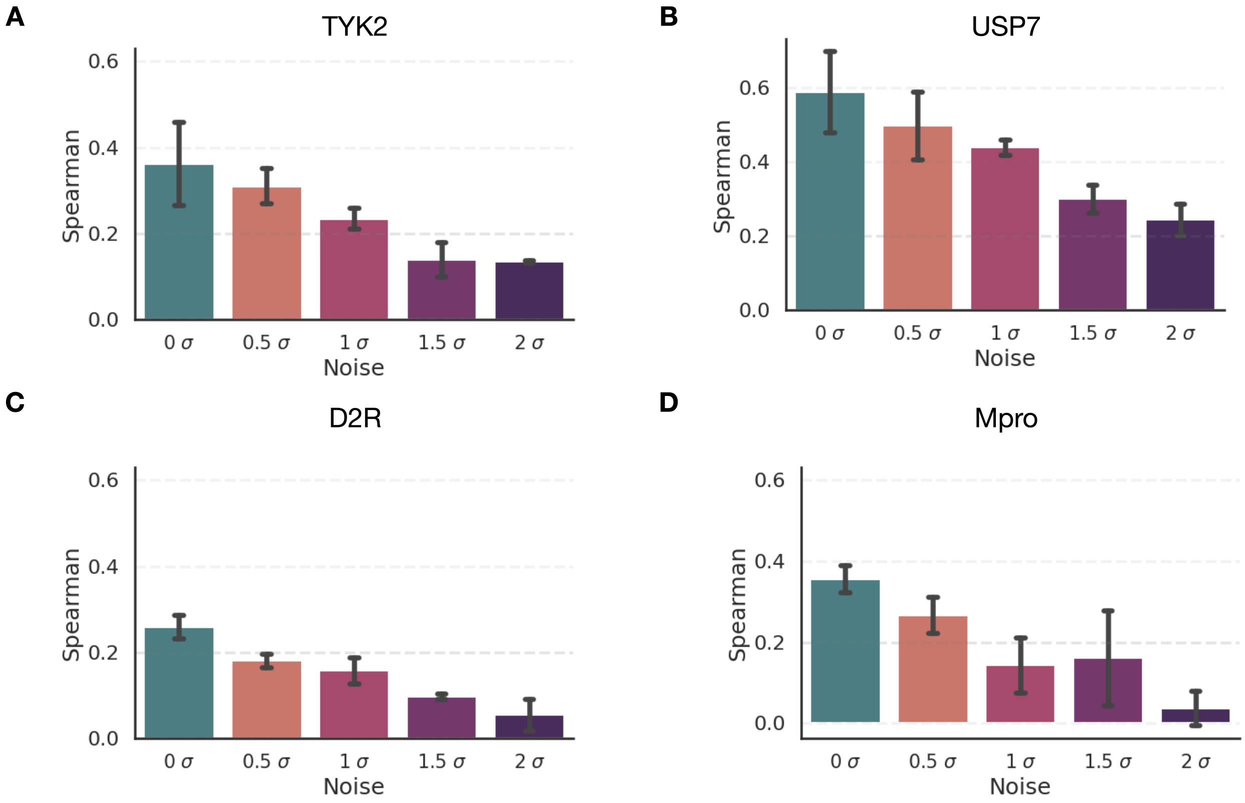


Figure S23: Spearman $\rho$ for CP models at different noise levels on for four datasets used in our study- **A:** TYK2, **B:** USP7, **C:** D2R, and **D:** Mpro.

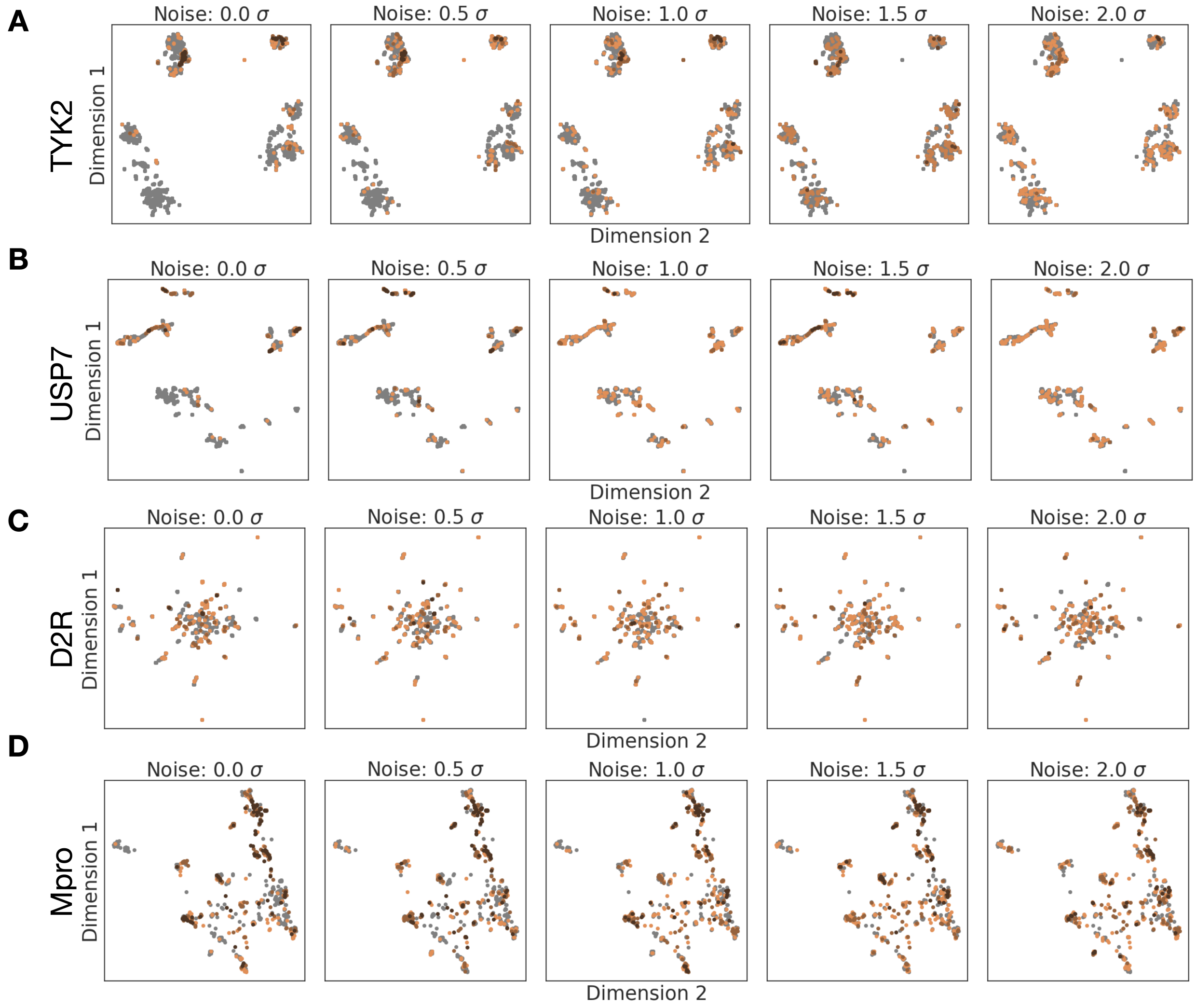


Figure S24: UMAPs using the CP model showing the chemical space changes at different noise levels on for four datasets used in our study- **A:** TYK2, **B:** USP7, **C:** D2R, and **D:** Mpro. UMAP visualization of the compounds selected in the exploitation phase, colored by the acquisition frequency across three distinct AL iterations with randomized initializations.

#### Selection of Initial Samples (on Training set)

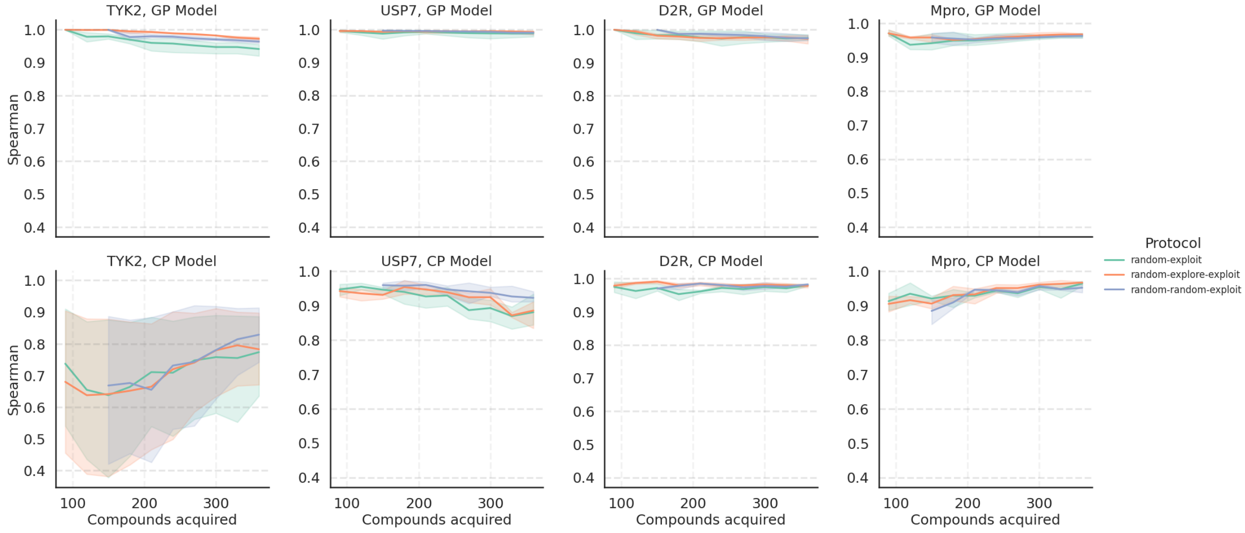


Figure S25: Spearman $\rho$ using different AL protocols on the training set (size equal to Compounds acquired) on all four target datasets with GP and CP models. Compounds acquired are cumulative over AL cycles. The shaded area is variation over 3 AL runs with different seeds.

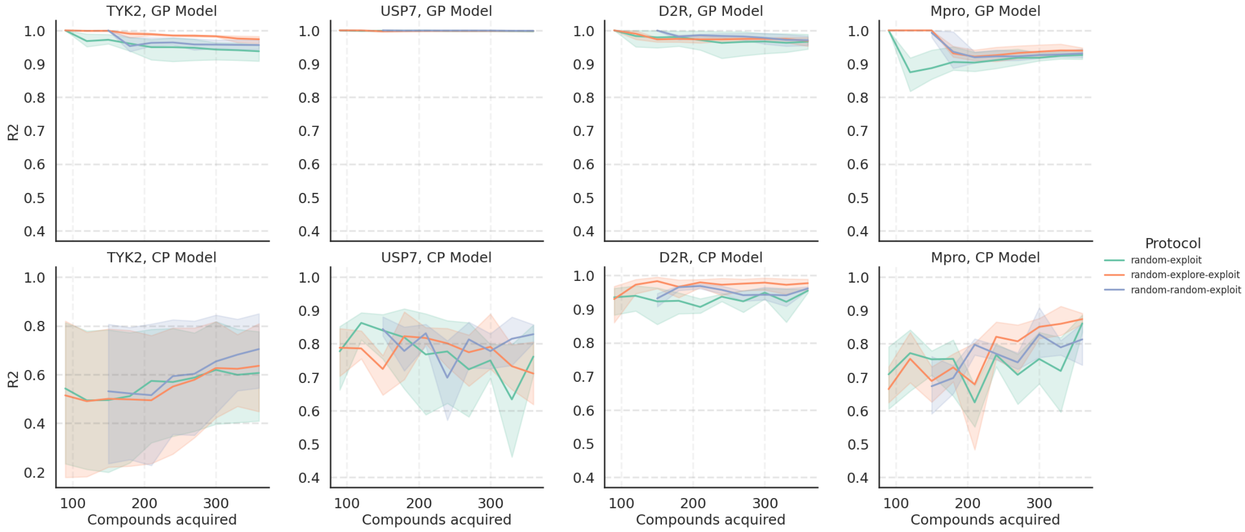


Figure S26: R2 on the training set (size equal to Compounds acquired) using different AL protocols on all four target datasets with GP and CP models. Compounds acquired are cumulative over AL cycles. The shaded area is variation over 3 AL runs with different seeds.

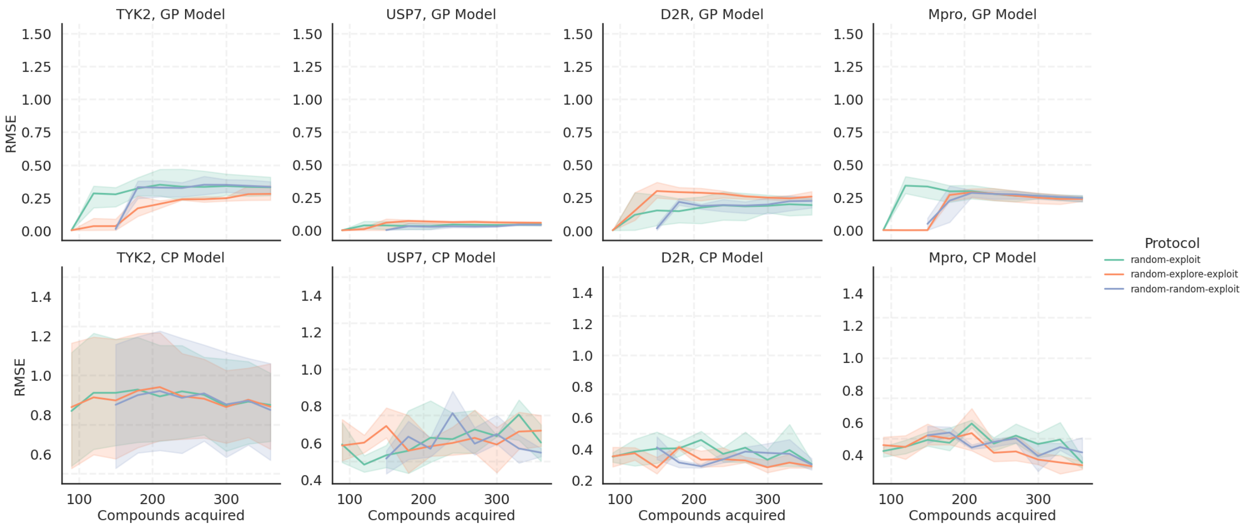


Figure S27: RMSE on the training set (size equal to Compounds acquired) using different AL protocols on all four target datasets with GP and CP models. Compounds acquired are cumulative over AL cycles. The shaded area is variation over 3 AL runs with different seeds.

#### Influence of batch size (on Training set)

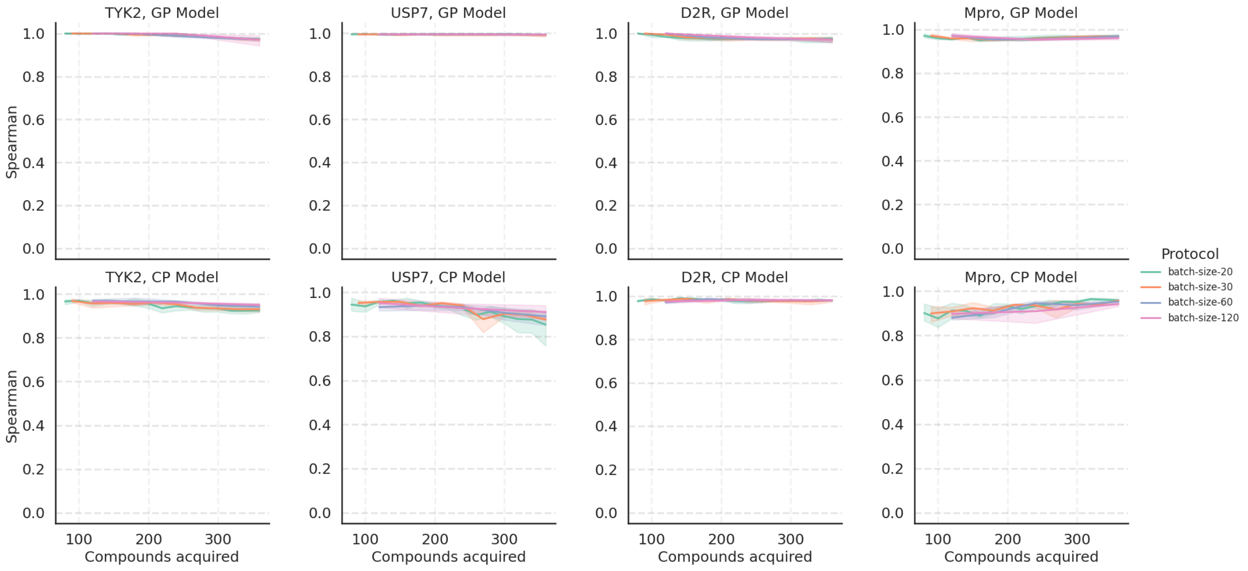


Figure S28: Spearman $\rho$ on the training set (size equal to Compounds acquired) using different AL protocols with varying batch sizes on all four target datasets with GP and CP models. Compounds acquired are cumulative over AL cycles. The shaded area is variation over 3 AL runs with different seeds.

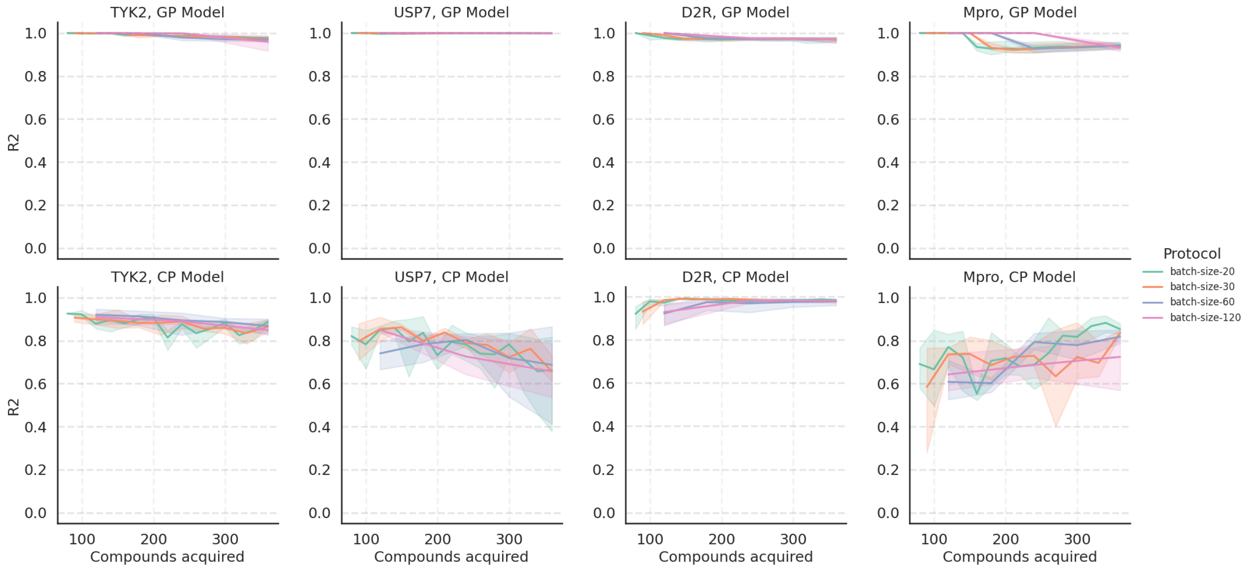


Figure S29: R2 on the training set (size equal to Compounds acquired) using different AL protocols with varying batch sizes on all four target datasets with GP and CP models. Compounds acquired are cumulative over AL cycles. The shaded area is variation over 3 AL runs with different seeds.

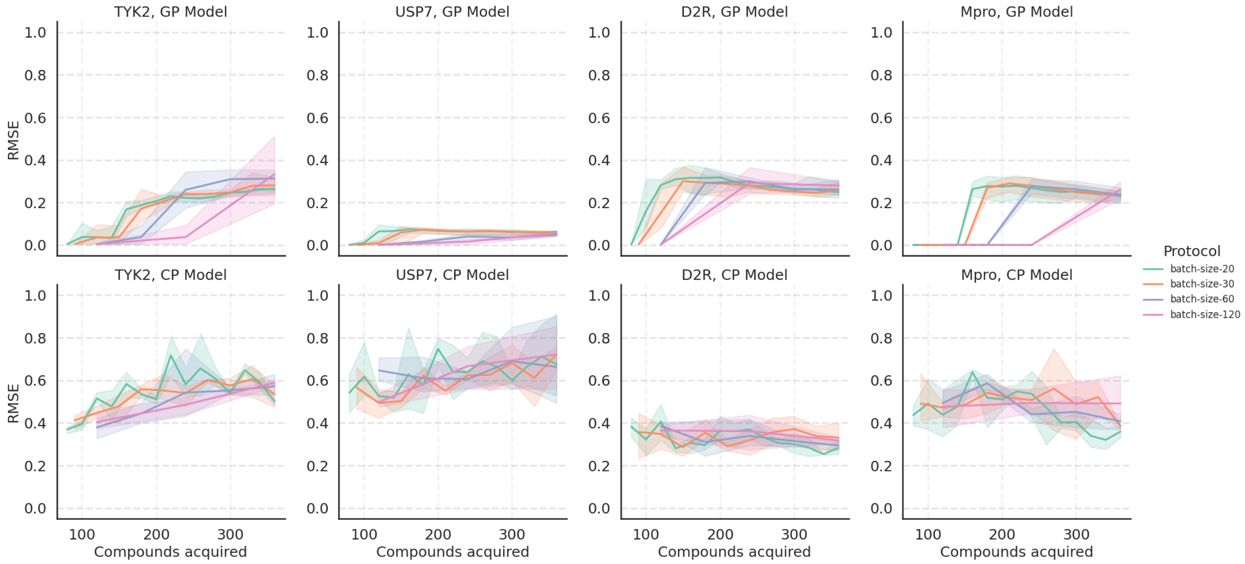


Figure S30: RMSE on the training set (size equal to Compounds acquired) using different AL protocols with varying batch sizes on all four target datasets with GP and CP models. Compounds acquired are cumulative over AL cycles. The shaded area is variation over 3 AL runs with different seeds.
